# Psilocybin triggers an activity-dependent rewiring of large-scale cortical networks

**DOI:** 10.1101/2025.08.06.668927

**Authors:** Quan Jiang, Lingxiao Shao, Shenqin Yao, Neil K. Savalia, Amelia D. Gilbert, Pasha A. Davoudian, Jack D. Nothnagel, Guilian Tian, Tin Shing Hung, Hei-Ming Lai, Kevin T. Beier, Hongkui Zeng, Alex C. Kwan

## Abstract

Psilocybin holds promise as a treatment for mental illnesses. One dose of psilocybin induces structural remodeling of dendritic spines in the medial frontal cortex in mice. The dendritic spines would be innervated by presynaptic neurons, but the sources of these inputs have not been identified. Here, using monosynaptic rabies tracing, we map the brain-wide distribution of inputs to frontal cortical pyramidal neurons. We discover that psilocybin’s effect on connectivity is network-specific: strengthening the routing of inputs from perceptual and medial regions (homolog of default mode network) to subcortical targets, while weakening inputs that are part of cortico-cortical recurrent loops. The pattern of synaptic reorganization depends on the drug-evoked spiking activity, because silencing a presynaptic region during psilocybin administration disrupts the rewiring. Collectively, the results reveal the impact of psilocybin on the connectivity of large-scale cortical networks and demonstrate neural activity modulation as an approach to sculpt the psychedelic-evoked neural plasticity.

## INTRODUCTION

Psychedelics are known to alter perceptual and cognitive states acutely after administration^1^. In the past decade, psychedelic-based treatments have emerged as a potential therapeutic for mental health conditions^2^. Among the various psychedelic compounds, psilocybin has yielded some of the most promising results. Numerous clinical trials have shown that psilocybin treatment with psychological support can relieve symptoms in patients with major depressive disorder or treatment-resistant depression^3–7^. Strikingly, the therapeutic benefit has been reported to last for at least 6 weeks after the administration of a single dose^5^. Psilocybin may be efficacious for other indications such as alcohol use disorder^8^. Because of the clinical relevance, there is an urgent need to understand the neurobiological mechanisms underlying psychedelic drug action^9^.

Structural neural plasticity is likely involved in psilocybin’s enduring effects on behavior. Individuals with major depressive disorder have fewer excitatory spine synapses and lower expression of synaptic proteins in the prefrontal cortex^10,11^. By contrast, the fast-acting antidepressant ketamine promotes the growth of dendritic spines in frontal cortical pyramidal neurons in rodents^12–14^. Like ketamine, a single dose of psilocybin leads to the formation of new dendritic spines in the mouse dorsal medial frontal cortex^15^. Of note, the elevation in spine number density induced by classic psychedelics is long-lasting, persisting for at least a month^15,16^. The durability of the structural remodeling in the brain may explain how psilocybin, which has a short half-life in the body^17^, can cause sustained changes in behavior.

Considering that dendritic spines are the postsynaptic sites for excitatory synapses, a key unanswered question is the identity of the presynaptic neurons that provide axonal inputs for the new connections formed after psilocybin administration. As part of the medial network of the mouse^18^, the dorsal medial frontal cortex receives long-range inputs from other cortical regions, such as the ventromedial prefrontal cortex, retrosplenial cortex, posterior parietal cortex, anterior insular cortex, as well as from subcortical regions, such as the mediodorsal and ventromedial nuclei of the thalamus, basolateral amygdala, and claustrum^18–20^. Additionally, there are local inputs and inter-hemispheric inputs from the contralateral hemisphere^19,20^. Uncovering the identity of the presynaptic neurons that provide inputs to the newly formed spines after drug administration is crucial, because it will reveal the specific neural pathways that are modified by psilocybin.

Here, we perform monosynaptic tracing to map the brain-wide distribution of presynaptic cells that send inputs to the mouse dorsal medial frontal cortex. We find that psilocybin administration alters neuronal connectivity in a manner that is highly specific to brain networks. The subcortical-projecting, pyramidal tract (PT) subtype of frontal cortical pyramidal neurons gain inputs almost exclusively from the medial, sensorimotor, and visual-auditory networks, but lose inputs from the ventromedial prefrontal cortex and lateral network. By contrast, the intratelencephalic (IT) subtype of frontal cortical pyramidal neurons exhibit a distinct and opposite pattern of reorganization for their presynaptic inputs. Moreover, we show that the acute effect of psilocybin on spiking activity determines if a particular source of presynaptic input would be strengthened subsequently. Collectively, our study provides crucial insights into how psilocybin modifies the connectivity of cortical networks.

## RESULTS

### Whole-brain tracing of drug-evoked difference in monosynaptic inputs

Monosynaptic tracing can be achieved using a rabies virus that is engineered to be EnvA-pseudotyped and glycoprotein (G) deficient^21^. Pseudotyping ensures that the rabies virus can enter only a set of experimenter-specified neurons (‘starter cells’) that express the EnvA receptor TVA. The deletion of G means that the rabies virus cannot spread, except from the starter cells in which G is expressed as a transgene. Therefore, a pseudotyped, G-deleted rabies virus (henceforth referred to as simply ‘rabies virus’) can transduce the starter cells, spread by crossing retrogradely one synapse to the presynaptic neurons (‘input cells’), but would halt there from further spread. In this study, we used two viruses for monosynaptic tracing: a Cre-dependent AAV helper virus, AAV-hSyn-DIO-TVA^66T^-dTomato-CVS N2c G, and the rabies virus, EnvA-CVS N2c^ΔG^-H2B-EGFP (**Figure 1A**). The viruses were engineered to have several favorable characteristics^22^. One, the TVA has a point mutation Glu^66^ to Thr to reduce EnvA-enveloped viral transduction, which mitigates the issue of non-specific, Cre-independent transduction of rabies virus due to leaky TVA expression^23^. Two, EGFP is fused to the histone H2B to restrict expression within the nucleus, enabling automated cell counting using machine learning methods. Three, the CVS N2c strain of rabies virus allows for efficient retrograde spread from cortical neurons, labeling more input cells per starter cell^24^.

**Figure 1.**
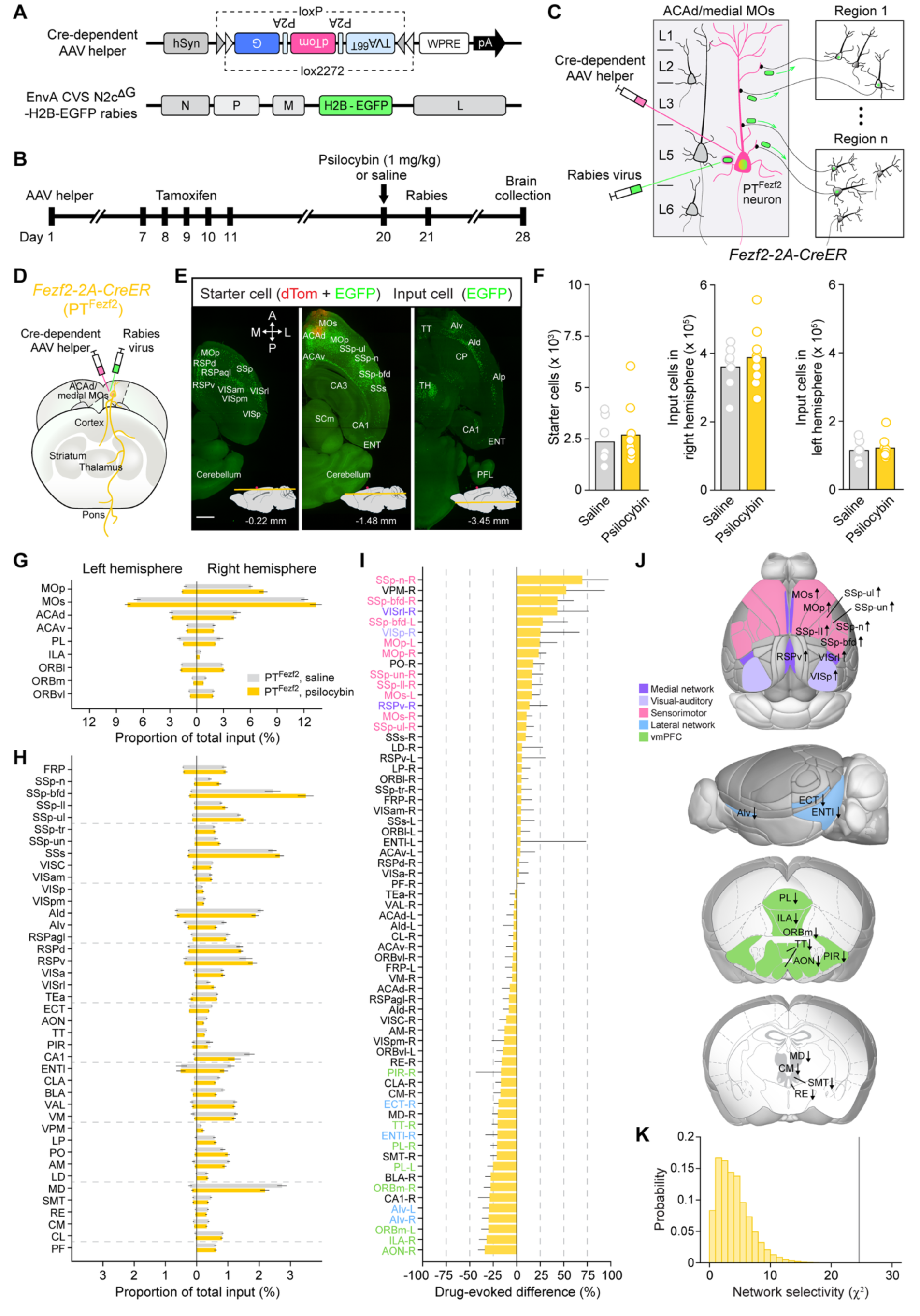
Psilocybin modifies inputs to frontal cortical PT^Fezf2^ neurons in a network-specific pattern. **(A)** The Cre-dependent AAV helper and the pseudotyped, G-deleted rabies virus used in this study. **(B)** Experimental timeline. **(C)** When the viruses are injected in the dorsal medial frontal cortex of a *Fezf2-2A-CreER* mouse, then the starter cells would be frontal cortical PT^Fezf2^ neurons. Starter cells would express dTomato and EGFP, while input cells would express EGFP only. **(D)** Monosynaptic tracing starting from the frontal cortical PT^Fezf2^ neurons using the *Fezf2-2A-CreER* mouse. **(E)** Three images in the horizontal plane from the whole-brain imaging, showing the right hemisphere of a *Fezf2-2A-CreER* mouse brain with starter cells (red and green) and input cells (green). The approximate depths relative to bregma (red circle) are indicated. Scale bar, 1 mm. **(F)** The total number of starter cells, input cells in the ipsilateral hemisphere, and input cells in the contralateral hemisphere for *Fezf2-CreER* mice treated with saline or psilocybin. Bar, mean. Circle, individual animal. Wilcoxon rank-sum test. **(G)** The proportion of input cells contributed by presynaptic regions in the frontal cortex in the left and right hemispheres (mean ± SEM). **(H)** The proportion of input cells contributed by other presynaptic regions in the left and right hemispheres (mean ± SEM). **(I)** Drug-evoked difference (psilocybin subtracted by saline, relative to saline) in the proportion of input cells for all 65 presynaptic regions for PT^Fezf2^ neurons (mean and 90% confidence interval). The list of presynaptic regions was sorted based on the drug-evoked difference value for PT^Fezf2^ neurons. Presynaptic regions with >10% or <-15% in drug-evoked difference were color-coded according to their network membership. **(J)** Schematic showing the location of the presynaptic regions with >10% or <-15% in drug-evoked difference for PT^Fezf2^ neurons. The presynaptic regions are color-coded according to their network membership. **(K)** Network selectivity analysis, testing against the null hypothesis that the increases and decreases in drug-evoked difference for PT^Fezf2^ neurons would be distributed randomly across the 5 cortical networks. Histogram, the distribution of χ^2^ values for the null hypothesis. Vertical line, the observed χ^2^ value (*P* = 6 × 10^-5^). ACAd, anterior cingulate area, dorsal part; ACAv, anterior cingulate area, ventral part; AId, agranular insular area, dorsal part; AIp, agranular insular area, posterior part; AIv, agranular insular area, ventral part; AON; anterior olfactory nucleus; CA1, field CA1; CA3, field CA3; CM, central medial nucleus of the thalamus; CP, caudoputamen; ECT, ectorhinal area; ENT, entorhinal area; MD, mediodorsal nucleus of the thalamus; MOp, primary motor area; MOs, secondary motor area; ORBl, orbital area, lateral part; ORBvl, orbital area, ventrolateral part; PFL, paraflocculus; PIR, piriform area; PL, prelimbic area; RE: nucleus of reuniens; RSPagl, retrosplenial area, lateral agranular part; RSPd, retrosplenial area, dorsal part; RSPv, retrosplenial area, ventral part; SCm, superior colliculus, motor related; SMT, submedial nucleus of the thalamus, SSp, primary somatosensory area; SSp-bfd, primary somatosensory area, barrel field; SSp-n, primary somatosensory area, nose; SSp-ul, primary somatosensory area, upper limb; SSs, supplemental somatosensory area; TH, thalamus; TT, taenia tecta; VISa, anterior area; VISam, anteromedial visual area; VISC, visceral area; VISp, primary visual area; VISpm, posteromedial visual area; VISrl, rostrolateral visual area; VPM, ventral posteromedial nucleus. N = 9 mice for psilocybin and 8 mice for saline.

The experimental timeline for monosynaptic tracing is shown in **Figure 1B**. The Cre-dependent AAV helper virus was injected into the dorsal medial frontal cortex of an animal, for example the *Fezf2-2A-CreER* mouse which is an inducible Cre-driver strain targeting the PT subtype of cortical pyramidal cells^25^. Subsequently, tamoxifen was administered to induce Cre expression. Two weeks later, after sufficient time for TVA and G expression in the starter cells, psilocybin (1 mg/kg, i.p.) or saline was administered, followed a day later by the injection of rabies virus into the dorsal medial frontal cortex. The rabies virus was injected after drug administration to ensure that the labelled input cells would reflect differences caused by psilocybin and not pre-existing connectivity prior to drug administration. We chose a dose of 1 mg/kg for psilocybin because it evokes structural neural plasticity in the mouse dorsal medial frontal cortex^15^. Finally, one week would elapse for rabies to spread before the brain was collected for analysis. The anticipated result is that we would find starter cells expressing both dTomato and EGFP, whereas the input cells would express EGFP only (**Figure 1C**).

We targeted the viral injections to the dorsal medial frontal cortex, encompassing the anterior cingulate cortex (ACAd) and the premotor cortex (medial portion of MOs), because brain-wide c-Fos mapping studies showed that this region responds robustly to stress, ketamine, and psilocybin^26,27^. Moreover, psilocybin leads to rapid and persistent structural neural plasticity in this region of the medial frontal cortex^15^. We verified that the monosynaptic tracing method worked as expected by performing multiple control experiments, involving a modified AAV helper virus that does not express G protein (**Figure S1A-C**), AAV helper virus only without rabies (**Figure S1D-E**), and rabies virus only without AAV helper virus (**Figure S1F-G**).

### Psilocybin modifies inputs to frontal cortical PT^Fezf2^ neurons in a network-specific pattern

Two of the major subtypes of pyramidal neurons in the neocortex are the PT and IT neurons (**Figure S2A-F**). These subtypes are non-overlapping populations with distinct morphological characteristics, electrophysiological properties, and long-range projection targets^19,28,29^.

Previous work indicated that the PT subtype of pyramidal neurons in the dorsal medial frontal cortex is essential for the long-term behavioral effects of psilocybin^30^. To determine how psilocybin may alter inputs into frontal cortical PT neurons, we performed monosynaptic tracing using *Fezf2-2A-CreER* mice (n = 4 male and 5 female animals for psilocybin, 4 male and 4 female animals for saline; **Figure 1D, S2G-H**). We injected viruses into the right hemisphere; therefore, the right hemisphere is ipsilateral and left hemisphere is contralateral. To determine the brain-wide distribution of input cells, we processed the rabies-traced brains through a pipeline involving tissue clearing, light sheet fluorescence microscopy, and machine learning-based automated detection of nuclei, which enabled us to locate and count all starter and input cells in the mouse brain (**Figure S3A**). As expected, we observed co-expression of red (dTomato) and green (EGFP) fluorescence from starter cells located in the dorsal medial frontal cortex, whereas green (EGFP) fluorescence from input cells was widespread in the brain (**Figure 1E**). **Video S1** shows images of the dTomato (red) and EGFP (green) expression, highlighting the brain-wide distribution of input cells into the frontal cortical PT^Fezf2^ neurons. **Video S2** shows images from the same brain for the dTomato expression (red) and NeuN immunostaining (white), showing the starter PT^Fezf2^ neurons with cell bodies in the dorsal medial frontal cortex and axons projecting ipsilaterally to the striatum as well as out of the cerebrum into the thalamus and pons. The images were aligned to the mouse brain atlas and divided based on the 316 summary structures in the Allen Mouse Brain Common Coordinate Framework version 3^31^.

Starter cells were defined as those neurons with colocalized red and green fluorescence within the ipsilateral frontal cortex (MOp, MOs, ACAd, ACAv, PL, ILA, ORBl, ORBm, and ORBvl). We had similar initial conditions for the psilocybin and saline groups, with 2680±491 and 2350±359 starter cells respectively (*P* = 0.7, Wilcoxon rank-sum test; **Figure 1F**). As expected, most starter cells were detected in the dorsal medial frontal cortex (94±2% and 95±2% in ACAd, MOs, or PL for the saline and psilocybin groups respectively; **Figure S3F**). Input cells were defined as those neurons with green fluorescence anywhere in the brain, except they cannot also have red fluorescence which would make them starter cells. For mice that received psilocybin, there were 5.1±0.5×10^5^ input cells, whereas for mice that received saline, there were 4.7±0.3×10^5^ input cells (*P* = 0.7, Wilcoxon rank-sum test; **Figure 1F, S3B, S3D**).

To identify the sources of the presynaptic inputs, we calculated the fraction of input cells residing in each region. Many regions did not provide appreciable input to the dorsal medial frontal cortex, so we analyzed only the 65 regions contributing at least 0.3% of the total inputs in any treatment or cell-type condition (henceforth referred to as ‘presynaptic regions’). Together these presynaptic regions captured nearly all input cells (saline: 87.6±0.6%; psilocybin: 89.2±0.7%; **Figure 1G, H**). Next, we sorted the presynaptic regions based on the normalized difference of psilocybin’s effect on the input fraction relative to saline, revealing numerous regions with increases, including the primary somatosensory cortex and its associated thalamic nuclei (SSp, VPM, PO), posterior parietal cortex (VISrl), primary visual cortex (VISp), primary motor cortex (MOp), retrosplenial cortex (RSP), and others (**Figure 1I**). Other regions had decreased input fraction following psilocybin administration, such as infralimbic area (ILA), medial orbital frontal cortex (ORBm), ventral agranular insular cortex (AIv), hippocampus (CA1), and basolateral amygdala (BLA). Focusing on the regions with largest psilocybin-evoked increase (>10%) or decrease (<-15%) in input fraction, we discovered that the strengthened presynaptic regions belong to the medial, visual-auditory, and sensorimotor networks (**Figure 1J**). This demarcation of cortical networks is based on modules identified in a prior large-scale study of long-range connectivity^18^. Regions that contributed fewer inputs after psilocybin administration came from the lateral network, ventromedial prefrontal cortex, and medial nuclei of the thalamus (**Figure 1J**). To test the statistical likelihood of such a network-specific preference in the input fraction change, we performed a chi-squared test to compare the observed data with shuffled data to find that the network selectivity for psilocybin’s effect was highly significant for PT^Fezf2^ neurons (χ^2^ = 24.6, d.f. = 4, *P* = 6 × 10^-5^, chi-squared test; **Figure 1K**). The results demonstrate that psilocybin induces a network-specific reorganization of the inputs impinging on the PT subtype of pyramidal neurons in the mouse dorsal medial frontal cortex.

### Opposing effects of psilocybin on inputs to the two major pyramidal cell types

Another major subtype of pyramidal neurons is the IT neurons. Unlike the subcortical-projecting PT neurons, the axons of IT neurons stay within the telencephalon to communicate with cortical and striatal locations in both hemispheres. Studies that examined cortical PT and IT neurons showed that the two cell types receive different local and long-range inputs^19,32–34^, which may underpin their distinct roles during behavior^30,35–37^. To determine how psilocybin may modify the inputs to frontal cortical IT neurons, we performed monosynaptic tracing using the *PlexinD1-2A-CreER* mouse, which is an inducible Cre-driver line for IT neurons^25^ (n = 4 male and 4 female animals for psilocybin, 4 male and 4 female animals for saline; **Figure 2A-B, S2G-H, Video S3**, **Video S4**).

**Figure 2.**
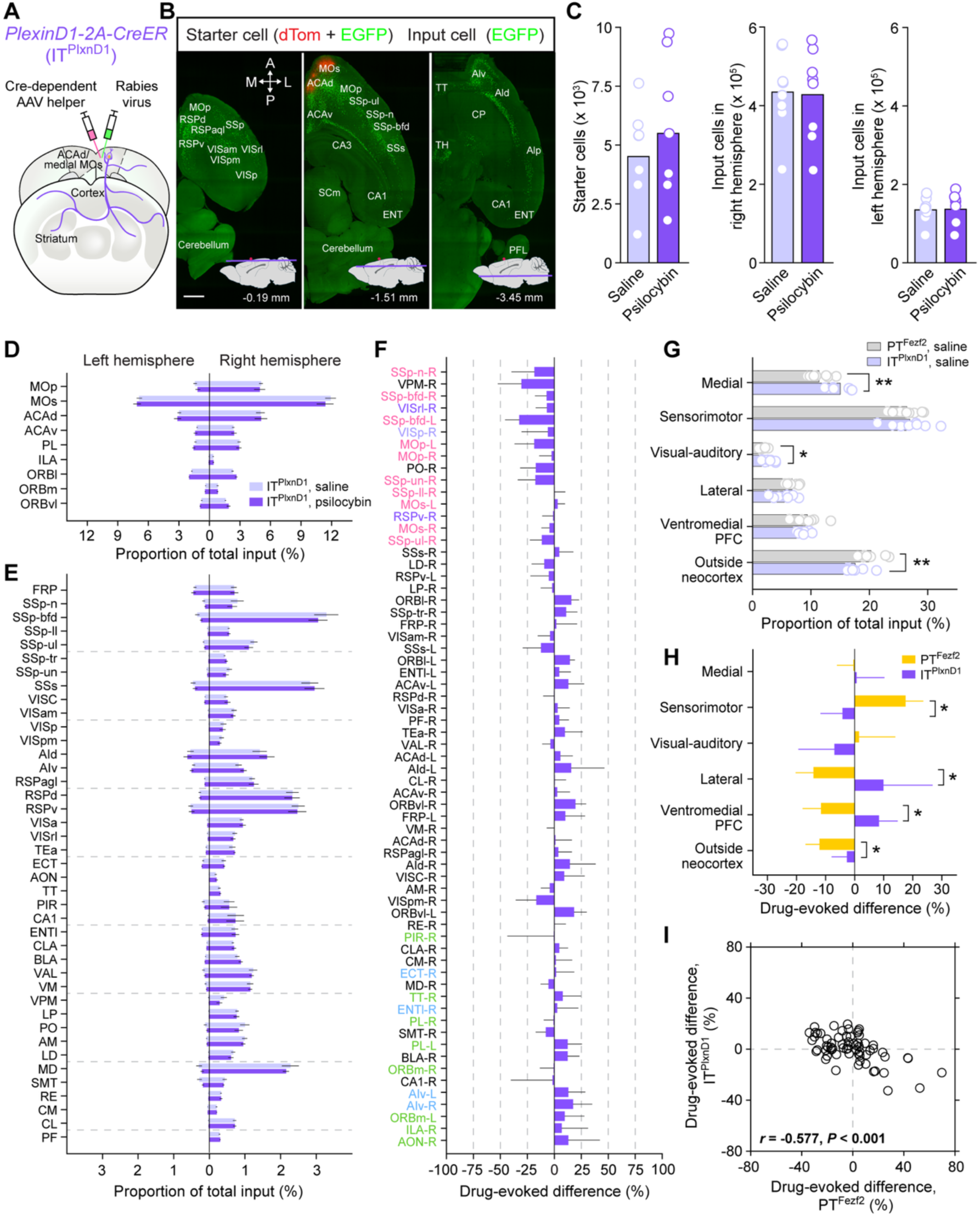
Opposing effects of psilocybin on presynaptic inputs to the two major pyramidal cell types. **(A)** Monosynaptic tracing starting from the frontal cortical IT^PlxnD1^ neurons using the *PlexinD1-2A-CreER* mouse. **(B)** Three images in the horizontal plane from the whole-brain imaging, showing the right hemisphere of a *PlexinD1-2A-CreER* mouse brain with starter cells (red and green) and input cells (green). The approximate depths relative to bregma (red circle) are indicated. Scale bar, 1 mm. **(C)** The total number of starter cells, input cells in the ipsilateral hemisphere, and input cells in the contralateral hemisphere for *PlexinD1-2A-CreER* mice treated with saline or psilocybin. Bar, mean. Circle, individual animal. Wilcoxon rank-sum test. **(D)** The proportion of input cells contributed by presynaptic regions in the frontal cortex in the left and right hemispheres (mean ± SEM). **(E)** The proportion of input cells contributed by other presynaptic regions in the left and right hemispheres (mean ± SEM). **(F)** Drug-evoked difference (psilocybin subtracted by saline, relative to saline) in the proportion of input cells for all 65 presynaptic regions for IT^PlxnD1^ neurons (mean and 90% confidence interval). The list of presynaptic regions was sorted based on the drug-evoked difference value for PT^Fezf2^ neurons (i.e., same order as Fig. 2F) and color-coded according to their network membership. **(G)** Source of ipsilateral presynaptic inputs to PT^Fezf2^ and IT^PlxnD1^ neurons in saline-treated mice. Circle, individual animal. *, *P* < 0.05, **, *P* < 0.01, Wilcoxon rank sum test. **(H)** Drug-evoked difference in the proportion of ipsilateral input cells from the 5 cortical networks and outside neocortex to PT^Fezf2^ and IT^PlxnD1^ neurons. Error bar, 90% confidence interval. *, the cell-type difference was significant based on 95% confidence interval. **(I)** Drug-evoked difference in the proportion of input cells for all 65 presynaptic regions, plotting results from tracing PT^Fezf2^ neurons versus IT^PlxnD1^ neurons. Circle, individual presynaptic region. *r*, Pearson correlation coefficient. N = 8 mice for psilocybin and 8 mice for saline. See also **Figure S2, S3, and S4**.

Beginning from 5495±1046 and 4511±696 starter cells (*P* = 0.6, Wilcoxon rank-sum test; 92±1% and 91±3% located in ACAd, MOs, or PL, **Figure 2C, S3G**), we counted 5.6±0.5×10^5^ and 5.7±0.5×10^5^ input cells for the psilocybin and saline groups respectively (*P* = 0.9, Wilcoxon rank-sum test; **Figure 2C, S3C, S3E**). The same set of 65 presynaptic regions, selected based on the criterion of containing at least 0.3% of the total inputs in any treatment or PT/IT condition, captured nearly all the input cells to the frontal cortical IT^PlxnD1^ neurons (saline: 89.4±0.6%; psilocybin: 89.8±0.6%). We plotted the input fraction for the psilocybin and saline groups in each of the frontal cortical (**Figure 2D**) and long-range cortical and subcortical presynaptic regions (**Figure 2E**). We plotted the difference in input fraction due to psilocybin relative to saline (**Figure 2F, S4A, S4D**). Although the specific input cells exhibiting changes were distinct between PT^Fezf2^ and IT^PlxnD1^ neurons, we found that psilocybin’s effect on the input fraction of frontal cortical IT^PlxnD1^ neurons is also highly network-specific (χ^2^ = 15.0, d.f. = 4, *P* = 0.005, chi-squared test; **Figure S4B-C**). The next analyses would be aimed at delineating the significant differences between PT^Fezf2^ and IT^PlxnD1^ neurons.

First, only considering the saline condition, we observed that PT^Fezf2^ and IT^PlxnD1^ neurons receive long-range inputs from different networks. Frontal cortical IT^PlxnD1^ neurons received a higher fraction of inputs from the medial network (15.0±0.7% for IT, 11.4±0.6% for PT; *P* = 0.004, Wilcoxon rank-sum test) and visual-auditory regions (2.7±0.3% for IT, 1.9±0.2% for PT; *P* = 0.05, Wilcoxon rank-sum test), but had significantly lower fraction of inputs from outside the neocortex (17.6±0.6% for IT, 20.3±0.7% for PT; *P* = 0.007, Wilcoxon rank-sum test), compared to their PT^Fezf2^ counterparts (**Figure 2G, S3H-I**). When we determined the effects of psilocybin, there were also cell-type-specific differences. In response to psilocybin, the IT^PlxnD1^ neurons differed significantly from the PT^Fezf2^ neurons in that they had reduced input fraction from the sensorimotor network (-4% for IT, 17% for PT; statistically significant based on 95% confidence interval from bootstrapping), but instead gained input fraction from lateral network (10% for IT, -14% for PT) and the ventromedial prefrontal cortex (8% for IT, -12% for PT; **Figure 2H**). Both pyramidal cell types lost input fraction from regions outside the neocortex after psilocybin administration, but the effect was less pronounced for IT neurons (-3% for IT, -12% for PT). Psilocybin’s opposing effect on the input fraction to the two cell types was most clearly demonstrated by plotting the drug-induced difference in input fraction for all 65 presynaptic regions, which showed a significant negative correlation (*r* = -0.58, *P* = 5×10^-7^, Pearson correlation; **Figure 2I**). These results indicate that, in the mouse dorsal medial frontal cortex, for those inputs that are strengthened in PT neurons by psilocybin, they tend to be weakened in IT neurons after drug administration, and vice versa. Therefore, psilocybin exerts opposing changes to the synaptic input organization for the two main pyramidal cell subtypes in the mouse dorsal medial frontal cortex.

### Psilocybin-evoked rewiring is unrelated to axonal innervation pattern

We wanted to know why inputs from certain presynaptic regions are selectively strengthened by psilocybin. We considered three possible factors. One, the presynaptic regions send axon collaterals that terminate in different layers of the mouse dorsal medial frontal cortex. Depending on the cortical depth, the long-range axons can access different dendritic compartments of the pyramidal neurons (e.g., axons in layer 1 would synapse onto the distal apical dendritic tuft), which may dictate psilocybin’s plasticity potential. Two, serotonin (5-HT) receptors are important for psilocybin’s drug action. A high baseline expression of certain subtypes of 5-HT receptors may predispose a presynaptic region to be sensitive to psilocybin-induced rewiring. Three, the presynaptic regions may have different acute responses to psilocybin. The pattern of synaptic reorganization can depend on the drug-evoked spiking activity of the neurons in the presynaptic region.

To test the first possibility regarding the laminar distribution of axons, we leveraged the Allen Mouse Connectivity database, which contains images of fluorescent axons following viral injections at hundreds of locations in the mouse brain^38^. We selected representative experiments in which AAV was injected in each of the 65 presynaptic regions and analyzed the images of fluorescent axons in the dorsal medial frontal cortex to extract the axonal density as a function of cortical layer. Although we always chose experiments in which the injection site centers at the presynaptic region, unavoidably the injections were often made over a larger extent than just the presynaptic region, which might influence the analysis. **Figure S5A** and **S5B** show 8 presynaptic regions with either increased or decreased input fraction after psilocybin based on monosynaptic tracing. **Figure S5C** and **S5D** are the corresponding laminar profile of the axonal density. We tested if various aspects of the axonal distribution may relate to the psilocybin-induced difference in input cells, but did not find any significant relationship. For example, we checked if the drug-evoked change in connectivity to PT^Fezf2^ neurons may relate to the amount of axons (**Figure S5E**), the laminar position for peak axonal density (**Figure S5F**), the fraction of axons residing in layer 1 (**Figure S5G**) or the fraction of axons residing in layer 2/3 (**Figure S5H**). These analyses suggest that the laminar distribution of long-range axons in the dorsal medial frontal cortex is unlikely to explain why certain presynaptic regions are favored for potentiation by psilocybin.

### Psilocybin-evoked rewiring is unrelated to the distribution of serotonin receptors

Next, we wanted to know if the expression of 5-HT receptors may relate to the network-specific reorganization of presynaptic inputs induced by psilocybin. In the body, psilocybin metabolizes into psilocin, which has affinity in the nanomolar range to numerous 5-HT receptor subtypes^1^, including the 5-HT_1A_, 5-HT_2A_, and 5-HT_2C_ receptors that are highly expressed in the neocortex^39^. We took advantage of the *in situ* hybridization data in the Allen Brain Atlas^39^ (**Figure 3A**), which has been curated previously to examine regional expression levels in the mouse neocortex^40,41^. We correlated the expression levels of the *Htr1a*, *Htr2a*, and *Htr2c* transcripts, which encode the 5-HT_1A_, 5-HT_2A_, and 5-HT_2C_ receptors respectively, with the psilocybin-induced difference in input cells for presynaptic regions in the cortex (**Figure 3B, C**). There was no trend for *Htr1a* and *Htr2a*. There may be an effect for *Htr2c*: although its regional expression level in the mouse cortex was not significantly related for PT^Fezf2^ neurons (*P* = 0.08, Pearson correlation), there may be a positive relationship for IT^PlxnD1^ neurons (*P* = 0.02, Pearson correlation).

**Figure 3.**
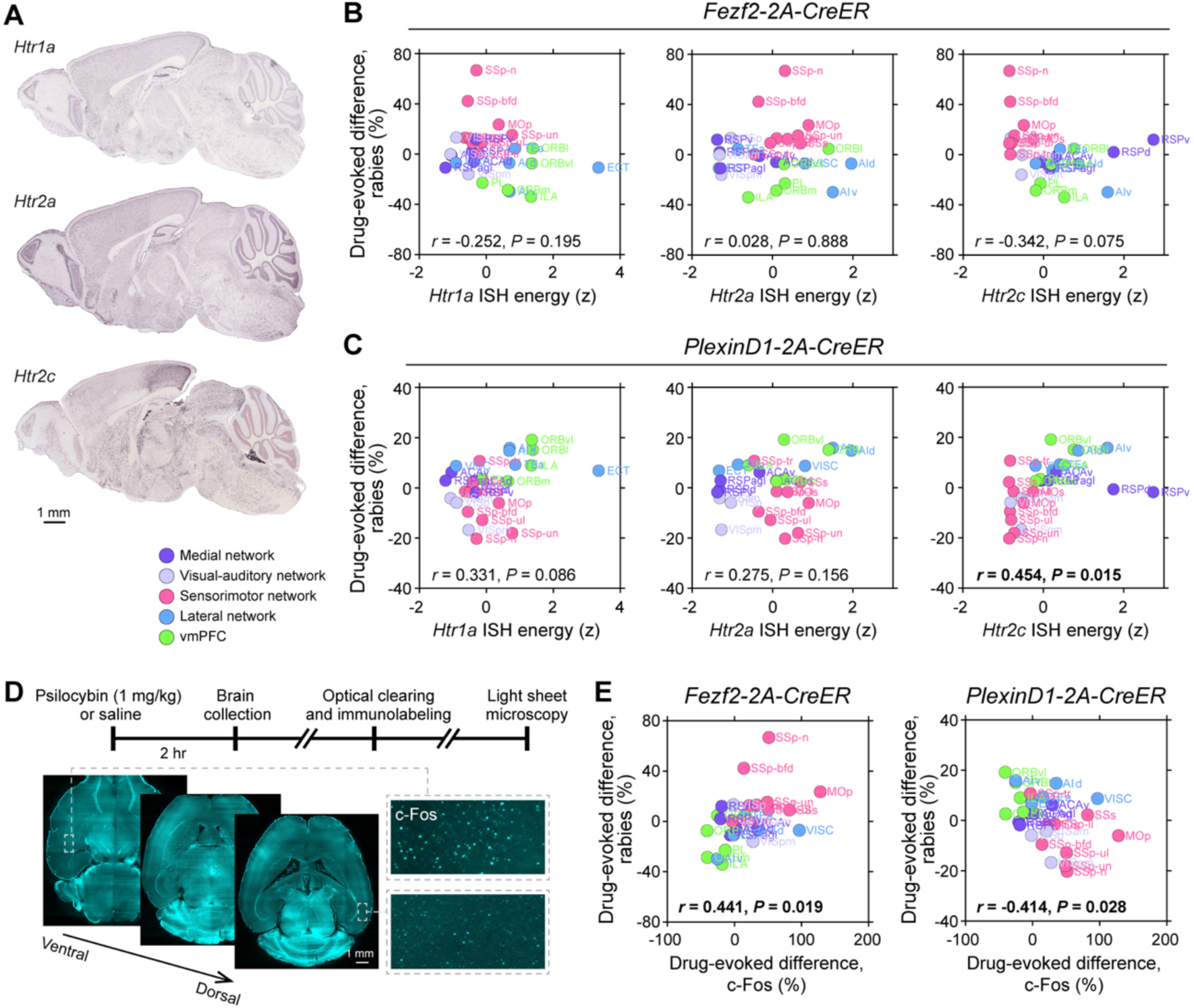
Psilocybin-evoked rewiring is unrelated to the distribution of serotonin receptors, but correlates with drug-evoked c-Fos expression. **(A)** *In situ* hybridization (ISH) images showing the expression of *Htr1a*, *Htr2a*, and *Htr2c* transcripts in the adult mouse brain. The images came from the Allen Institute for Brain Science database (http://www.brain-map.org). **(B)** The ISH expression of *Htr1a*, *Htr2a*, or *Htr2c* versus the psilocybin-evoked differences in input fraction for PT^Fezf2^ neurons. Circle, individual presynaptic region. The color denotes the cortical network membership. *r*, Pearson correlation coefficient. **(C)** Similar to (B) for IT^PlxnD1^ neurons. **(D)** Experimental timeline for imaging psilocybin’s effect on brain-wide c-Fos expression. Insets, representative fluorescence images in a psilocybin-treated mouse. **(E)** The psilocybin-evoked difference in c-Fos expression versus the psilocybin-evoked differences in input fraction for PT^Fezf2^ and IT^PlxnD1^ neurons. Circle, individual presynaptic region. The color denotes the cortical network membership. *r*, Pearson correlation coefficient. See also **Figure S5**.

We found a more robust relationship when examining the psilocybin-evoked activation of the immediate early gene c-Fos. In a previous study^42^, our lab has injected mice with psilocybin (1 mg/kg, i.p.) or saline, collected the brains 2 hours later, and then used tissue clearing, c-Fos antibody labeling, and light sheet fluorescence microscopy to visualize c-Fos+ cells in the entire mouse brain (**Figure 3D**). We compared the drug-evoked difference in c-Fos+ cells with the drug-evoked difference in input cells on a region-by-region basis. For the presynaptic regions in the neocortex, those locations with greater increases in the number of c-Fos+ cells were associated with a higher elevation of input cells for frontal cortical PT^Fezf2^ neurons after psilocybin administration (*r* = 0.44, *P* = 0.02, Pearson correlation; **Figure 3E**). By contrast, psilocybin’s effects on c-Fos activation and input cell gain were negatively correlated for the frontal cortical IT^PlxnD1^ neurons (*r* = -0.41, *P* = 0.03, Pearson correlation; **Figure 3E**). Altogether, these results indicate that psilocybin-induced c-Fos activation is related to the gain and loss of input cells for frontal cortical PT and IT neurons respectively. Because c-Fos expression is known to be activity-dependent, this exploratory analysis hints at spiking activity as a potential factor underlying the pattern of synaptic reorganization evoked by psilocybin.

### Potentiation of RSP inputs onto PT^Fezf2^ neurons in the medial frontal cortex after psilocybin administration

To delineate the potential mechanisms, we focused on one presynaptic region: the retrosplenial cortex (RSP). RSP is known to have dense reciprocal connections with the dorsal medial prefrontal cortex in mice^18,20,43^. RSP is relevant for the neurobiology of psilocybin, because it is a core region in the mouse analog of the default mode network^44–46^, which is thought to be centrally involved in mediating the effects of psychedelics in humans^47–49^. Our whole-brain rabies tracing results indicated that RSP was a top-ranked region with >10% increase in input fraction to frontal cortical PT^Fezf2^ neurons.

We performed longitudinal two-photon imaging to determine the effect of psilocybin on the density of long-range inputs in the medial frontal cortex (**Figure 4A, B**). To label the boutons along axons originating from neurons in RSP, we injected AAV-hSyn-Synaptophysin-mRuby2 into RSP. Initially, we planned to visualize simultaneously axonal boutons and dendritic spines, therefore also injected AAV-CAG-Flex-EGFP into ACAd/medial MOs of these *Fezf2-2A-CreER* mice to label PT^Fezf2^ neurons; however, the EGFP-expressing dendrites were too dense to resolve most spines and thus we restricted the analysis to axonal boutons only (**Figure 4C**). We tracked a total of 60 and 62 fields of view from 7 and 8 animals for the saline and psilocybin conditions, respectively. Using an automated algorithm to count red puncta from the *in vivo* images, we observed that psilocybin increased the number of detectable RSP axonal boutons in superficial layers of dorsal medial frontal cortex at 1 and 3 days following treatment, relative to saline (day 1: 5.5±3.3% for psilocybin versus -8.6±3.9% for saline; *P* = 0.056, post hoc comparison; day 3: 11.4±4.6% for psilocybin versus -7.6±3.9% for saline; *P* = 0.007, post hoc comparison; treatment and time interaction: *P* = 0.0004, linear mixed effects model; **Figure 4D**). We note a decline in bouton density prior to treatment, possibly due to photobleaching of the synaptophysin-fused fluorophores, which affected the saline and psilocybin groups similarly (day -1: -4.7±1.3% for psilocybin and -4.8±1.7% for saline; *P* = 1, post hoc comparison). The imaging results thus corroborate the finding from rabies tracing to show a psilocybin-evoked increase in RSP axonal boutons in the dorsal medial frontal cortex.

**Figure 4.**
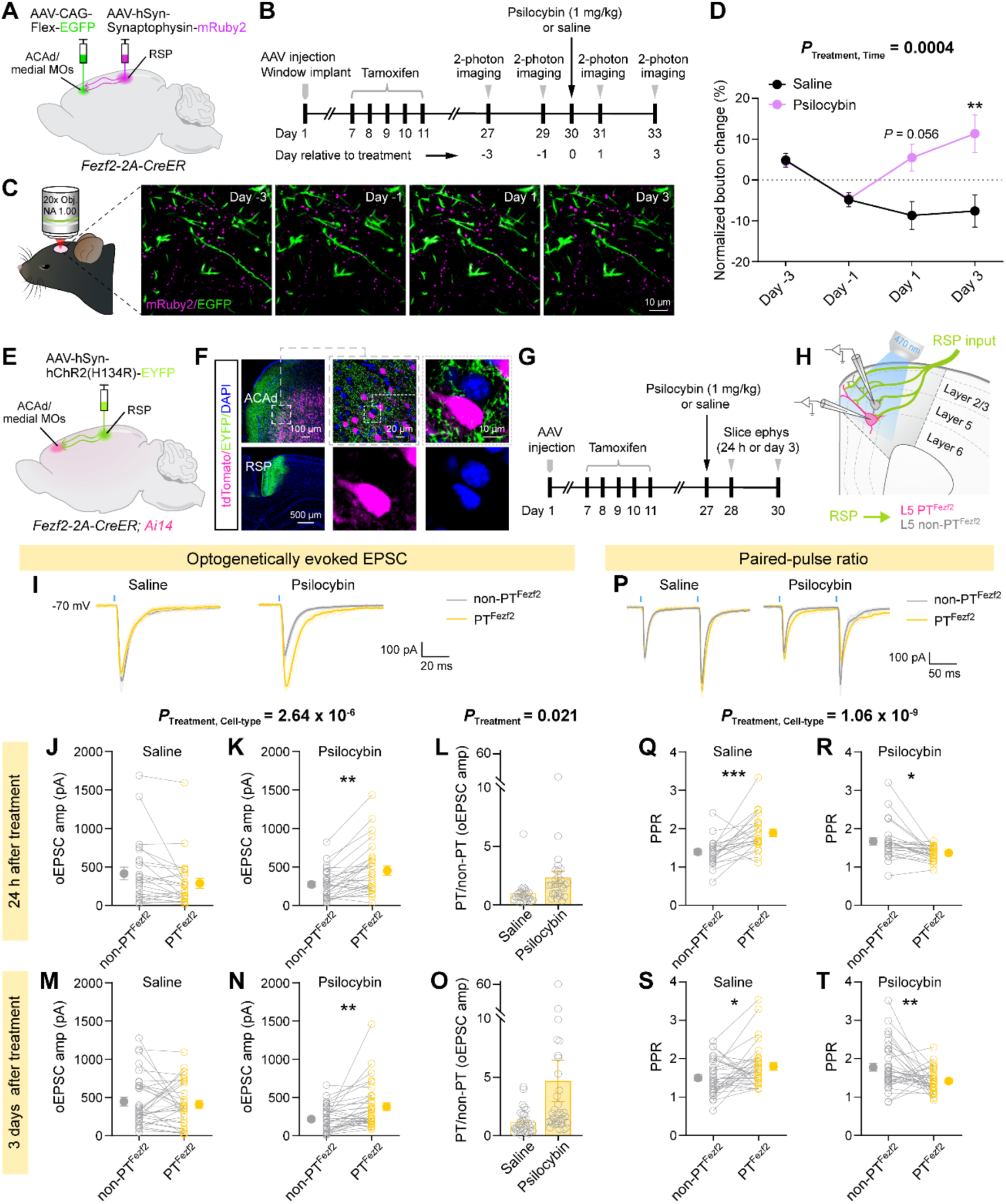
Potentiation of RSP inputs onto PT^Fezf2^ neurons in the medial frontal cortex after psilocybin administration. **(A)** Viral strategy to express mRuby2 in axonal boutons of RSP neurons and EGFP in frontal cortical PT^Fezf2^ neurons. **(B)** Experimental timeline. **(C)** Example *in vivo* two-photon images, tracking the same field of view at day -3, -1, 1, and 3 relative to psilocybin administration. **(D)** The density of RSP axonal boutons in the medial frontal cortex after treatment with psilocybin (magenta; 1 mg per kg, i.p.) or saline (black) across days, expressed as the fold change from the baseline defined as the mean value from the two pre-treatment sessions (day -3 and -1). **(E)** Viral strategy to express ChR2 in RSP neurons. PT^Fezf2^ neurons express tdTomato in the *Fezf2-2A-CreER;Ai14* mouse. **(F)** Post hoc histology showing the fluorophore expression in medial frontal cortex and RSP. **(G)** Experimental timeline. **(H)** Successive whole cell recordings were made from a PT^Fezf2^ neuron and a non-PT^Fezf2^ neuron. A 470 nm LED (2 ms) was used to photostimulate the RSP axons in the acute brain slice. **(I)** Example optogenetically evoked EPSC in a pair of PT^Fezf2^ and non-PT^Fezf2^ neurons, for saline and psilocybin condition. **(J)** Amplitude of the optogenetically evoked EPSC, 24 hr after saline administration. Circle, individual cell. **(K)** Similar to (J) for psilocybin administration. **(L)** Based on data in (J) and (K), the ratio calculated by dividing the amplitude of a PT^Fezf2^ neuron by the amplitude of its paired non-PT^Fezf2^ neuron. Circle, individual cell pair. Bar, mean ± SEM. **(M–O)** Similar to (J–L), 3 days after saline or psilocybin administration. **(P)** Example optogenetically evoked EPSCs evoked by two brief pulses in a pair of PT^Fezf2^ and non-PT^Fezf2^ neurons, for saline and psilocybin condition. **(Q)** Paired pulse ratio, 24 hr after saline administration. Circle, individual cell. **(R)** Similar to (Q) for psilocybin administration. **(S, T)** Similar to (Q, R), 3 days after saline or psilocybin administration. For imaging, N = 60 fields of view from 7 mice for saline and 62 fields of view from 8 mice for psilocybin. For slice electrophysiology, N = 23–25 cell pairs from 6 mice for 24 hr after saline, 26–29 cell pairs from 7 mice for 24 hr after psilocybin, 33 cell pairs from 5 mice for 3 days after saline, 34 cell pairs from 7 mice for 3 days after psilocybin. For (D), linear mixed effects model with fixed effects terms of drug (saline or psilocybin), time (-3, -1, 1, 3), and interaction, with mouse and field of view modeled as nested random intercepts. For (J, K, M, N) and (Q, R, S, T), linear mixed effects model with fixed effects terms of drug (saline or psilocybin), time (24 hr or 3 day), cell type (PT^Fezf2^ or non-PT^Fezf2^) and all interactions, with cell pairs per brain slice per mouse modeled as nested random intercepts. For (L, O), linear mixed effects model with fixed effects terms of drug (saline or psilocybin), time (24 hr or 3 day), and interaction, with random intercept for brain slice and animal. Post hoc pairwise comparisons with Bonferroni correction. See **Table S1**. *, *P* < 0.05. **, *P* < 0.01. ***, *P* < 0.001. See also **Figure S6**.

To evaluate further the effect of psilocybin on RSP inputs and specifically in frontal cortical PT^Fezf2^ neurons, we measured synaptic transmission using slice electrophysiology. We injected AAV-hSyn-hChR2(H134R)-EYFP into RSP to enable photostimulation of axons originating from RSP, which was done in *Fezf2-2A-CreER;Ai14* mice where PT^Fezf2^ neurons expressed tdTomato so we could target them for whole-cell recording (**Figure 4E, F**). Acute brain slices containing ACAd/medial MOs were prepared at 1 or 3 days after the administration of psilocybin (1 mg/kg, i.p.) or saline (5 - 7 mice for each condition; **Figure 4G**). During each recording session, we would pair the recordings by measuring in succession from a PT^Fezf2^ neuron and a non-PT^Fezf2^ neuron (i.e., an adjacent pyramidal cell without tdTomato) (**Figure 4H**). The motivation was to account for the variable ChR2 expression across experiments and the opposing effects of psilocybin for inputs onto PT^Fezf2^ and IT^PlxnD1^ neurons. A single brief pulse of photostimulation was used to elicit an optogenetically evoked excitatory postsynaptic current (oEPSC; **Figure 4I**). We found that psilocybin induced cell-type-specific effects on oEPSC amplitude when tested 24 hours or 3 days after treatment (treatment and cell type interaction: *P* = 3 x 10^-6^, linear mixed effects model; **Figure 4J, K, M, N**). Specifically, the oEPSC amplitude for PT^Fezf2^ neurons relative to non-PT^Fezf2^ pyramidal cells was higher after psilocybin administration (main effect of treatment: *P* = 0.02, linear mixed effects model; **Figure 4L, O**). We also assessed psilocybin’s effects on optogenetically evoked inhibitory postsynaptic currents (oIPSC; **Figure S6A-G**), the oEPSC-to-oIPSC ratio (**Figure S6H-J**), and monosynaptic oEPSC in the presence of TTX and 4-AP (**Figure S6K-O**). Two brief pulses of photostimulation spaced by a short interval was applied to measure paired-pulse ratio (PPR; **Figure 4P**). With this protocol, we likewise observed that psilocybin induced cell-type-specific effects on PPR at both 1 and 3 days after psilocybin administration (treatment and cell type interaction: *P* = 1 x 10^-9^, linear mixed effects model; **Figure 4Q-T**). The reduced PPR in PT^Fezf2^ neurons suggests psilocybin’s effects on the RSP→ACAd excitatory connections may include an increase in presynaptic release probability. Additional electrophysiological data obtained via a different viral labeling strategy provided further evidence for psilocybin’s effects on oEPSC and PPR (**Figure S6P-W**). Overall, these data demonstrate psilocybin-induced potentiation of excitatory synaptic drive from RSP to frontal cortical PT^Fezf2^ neurons, which persisted to at least 3 days after drug exposure.

### Psilocybin elevates the spiking activity of frontal cortex-projecting neurons in RSP

After confirming RSP as an input source altered by psilocybin, we tested the importance of spiking activity by measuring firing rates in RSP using *in vivo* electrophysiology. To record specifically from the frontal cortex-projecting neurons, we injected the retrogradely transported AAVretro-hSyn-Cre into the dorsal medial frontal cortex of an *Ai32* mouse for Cre-dependent expression of channelrhodopsin (ChR2) (**Figure 5A**). For each recording session, we would perform a craniotomy at ipsilateral RSP, insert a high-density Neuropixels electrode, record for 30 minutes, infuse psilocybin (1 mg/kg, i.p.) or saline through an implanted catheter, record for another 60 minutes, and then use laser photostimulation (473 nm, 20 ms) for opto-tagging (**Figure 5B**). We isolated single units from the recording via spike sorting and curated based on quality metrics (**Figure S7A-B**). The Neuropixels probe was coated with the fluorescent dye CM-DiI, allowing us to visualize the tract in post hoc histology performed after each recording session (**Figure 5C**). The reconstructed probe trajectories for all recordings confirmed the targeting of RSP (**Figure 5D**). During the opto-tag protocol, most neurons exhibited no detectable change in firing rate from the laser photostimulation, however some cells were driven to fire reliably and with short latency (**Figure 5E-F, S7C**). These neurons with photostimulation-evoked spiking activity would be the ChR2-expressing RSP→ACAd neurons.

**Figure 5.**
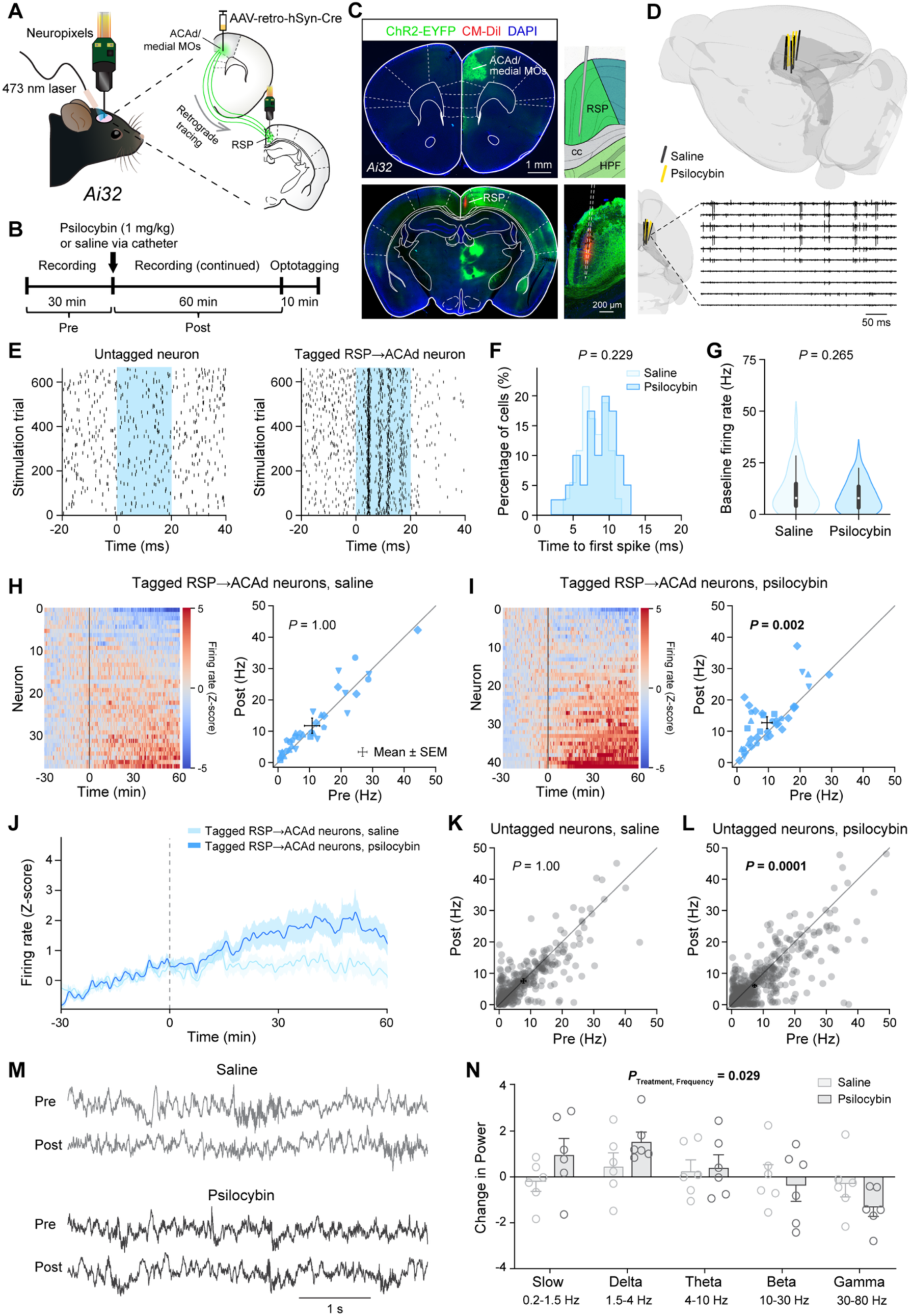
Psilocybin elevates the spiking activity of frontal cortex-projecting neurons in the retrosplenial cortex. **(A)** Experimental setup. **(B)** Experimental timeline. **(C)** Histology showing the injection site in the dorsal medial frontal cortex and the probe track in RSP. Inset, magnified view of the probe placement. **(D)** Probe tracks for all recordings reconstructed in the Allen Mouse Brain Common Coordinate Framework. Example segment of recording from 10 consecutive channels along the probe. **(E)** Spike raster plots for a non-tagged RSP neuron and a tagged RSP→ACAd neuron. Blue, period of laser stimulation. **(F)** The time of the first spike after laser onset for all tagged RSP→ACAd neurons in saline and psilocybin groups (KS statistic = 0.22, *P* = 0.2, two-sided two-sample Kolmogorov-Smirnov test). **(G)** Baseline firing rates of tagged RSP→ACAd neurons in saline and psilocybin groups, during the ‘Pre’ period. (*P* = 0.3, two-sample *t*-test). **(H)** Heatmaps showing z-scored firing rate for all tagged RSP→ACAd neurons before and after saline administration. Scatter plot of baseline (Pre) and post-saline (Post) firing rates for tagged RSP→ACAd neurons. Solid symbols represent individual neurons. Different symbol shapes indicate different animals. Crosshairs, mean ± SEM. Linear mixed effects model with fixed effects terms of treatment (saline or psilocybin), time (pre- or post-drug), and cell type (opto-tagged or untagged), and all interactions, as well as cell and mouse modeled as nested random intercepts. See **Table S1**. **(I)** Similar to (H) for psilocybin administration. **(J)** The z-scored firing rate as a function of time for tagged RSP→ACAd neurons in saline and psilocybin groups. Line, mean. Shading, SEM. **(K)** Similar to (H) for untagged neurons and saline administration. **(L)** Similar to (H) for untagged neurons and psilocybin administration. **(M)** Example LFP signals from the channel at a depth of 300 µm, in an animal before and after saline (top rows) or in another animal before and after psilocybin (bottom rows). **(N)** Fractional change in spectral power within different frequency after saline or psilocybin. Circle, individual animal. Bar, mean ± SEM. Linear mixed effects model with fixed effects terms of drug (saline or psilocybin), frequency (slow, delta, alpha, beta, or gamma), and interaction, with a random intercept for animal. See **Table S1**. N = 38 opto-tagged neurons from 6 mice for saline and 42 opto-tagged neurons from 6 mice for psilocybin. See also **Figure S7**.

In total, we recorded from 38 and 42 opto-tagged RSP→ACAd neurons, along with 417 and 710 other untagged neurons, in saline and psilocybin conditions respectively (n = 2 male and 4 female animals for psilocybin, 3 male and 3 female animals for saline). We compared the firing rates of opto-tagged RSP→ACAd neurons before and after the administration of psilocybin or saline (**Figure 5G-I, S7D-E**). For saline, there was no detectable change in the spiking activity of RSP→ACAd neurons (pre-saline: 11.1±1.6 Hz, post-saline: 12.2±1.6 Hz); by contrast, the administration of psilocybin elicited a 39% increase in the firing rate of the RSP→ACAd neurons (pre-psilocybin: 8.9±1.1 Hz, post-psilocybin: 12.4±1.3 Hz; *P* = 0.002, post hoc comparison; treatment, time, and cell type interaction: *P* = 0.031, linear mixed effects model). The psilocybin-induced elevation of spiking activity in RSP→ACAd neurons started at ∼15 minutes after drug administration and persisted until the end of the recording (**Figure 5J**). Psilocybin reduced the firing of untagged neurons (pre-saline: 7.8±0.5 Hz, post-saline: 7.6±0.5 Hz; *P* = 1, post hoc comparison; pre-psilocybin: 7.1±0.4 Hz, post-psilocybin: 6.1±0.3 Hz; *P* = 0.0001, post hoc comparison; **Figure 5K-L, S7F-G**). One caveat for this experiment is that the recorded RSP→ACAd neurons were not restricted to those that specifically project to PT^Fezf2^ or IT^PlxnD1^ cell types. In addition to single-unit activity, we analyzed the local field potential (LFP) recorded after drug administration (**Figures 5M**). Averaging the power spectra obtained from LFPs recorded from channels located within 300 µm of the pial surface and comparing pre-versus post-treatment, we observed a shift in the spectral power with increases around the delta band and decreases at the high-frequency gamma band after psilocybin administration (delta: 1.6±0.4% for psilocybin versus 0.5±0.6% for saline; gamma: -1.4±0.4% for psilocybin versus - 0.3±0.5% for saline; treatment and frequency interaction: *P* = 0.029, linear mixed effects model; **Figure 5N, S7H-L**). The increase in delta power was reminiscent of the oscillatory activity previously associated with the dissociative effects of ketamine^50,51^. Altogether, these data indicate that psilocybin increases spiking activity selectively in the frontal cortex-projecting neurons in RSP, which is one of the presynaptic regions that showed strengthened presynaptic inputs to the PT^Fezf2^ neurons after psilocybin.

### Chemogenetic silencing of RSP alters psilocybin-evoked rewiring

To test causally if the neural activity in a presynaptic region influences psilocybin-induced rewiring, we manipulated the excitability of neurons in RSP using chemogenetics^52^ (**Figure 6A**). We used the same experimental protocol as our monosynaptic tracing in the *Fezf2-2A-CreER* mouse, with the addition of injecting AAV-hSyn-hM4D(Gi)-mCherry in the RSP to express the inhibitory chemogenetic receptor hM4D(Gi) in RSP neurons (**Figure 6B**). At 15 min before the administration of psilocybin (1 mg/kg, i.p.) or saline, we would inject the chemogenetic ligand deschloroclozapine^53^ (DCZ; 0.1 mg/kg, i.p.) or DMSO vehicle. Therefore, we were using chemogenetics to silence RSP neurons when psilocybin is psychoactive. The experiment proceeded with rabies tracing and histology where coronal brain sections were prepared for visualization of EGFP-expressing input cells in RSP and neighboring brain regions.

**Figure 6.**
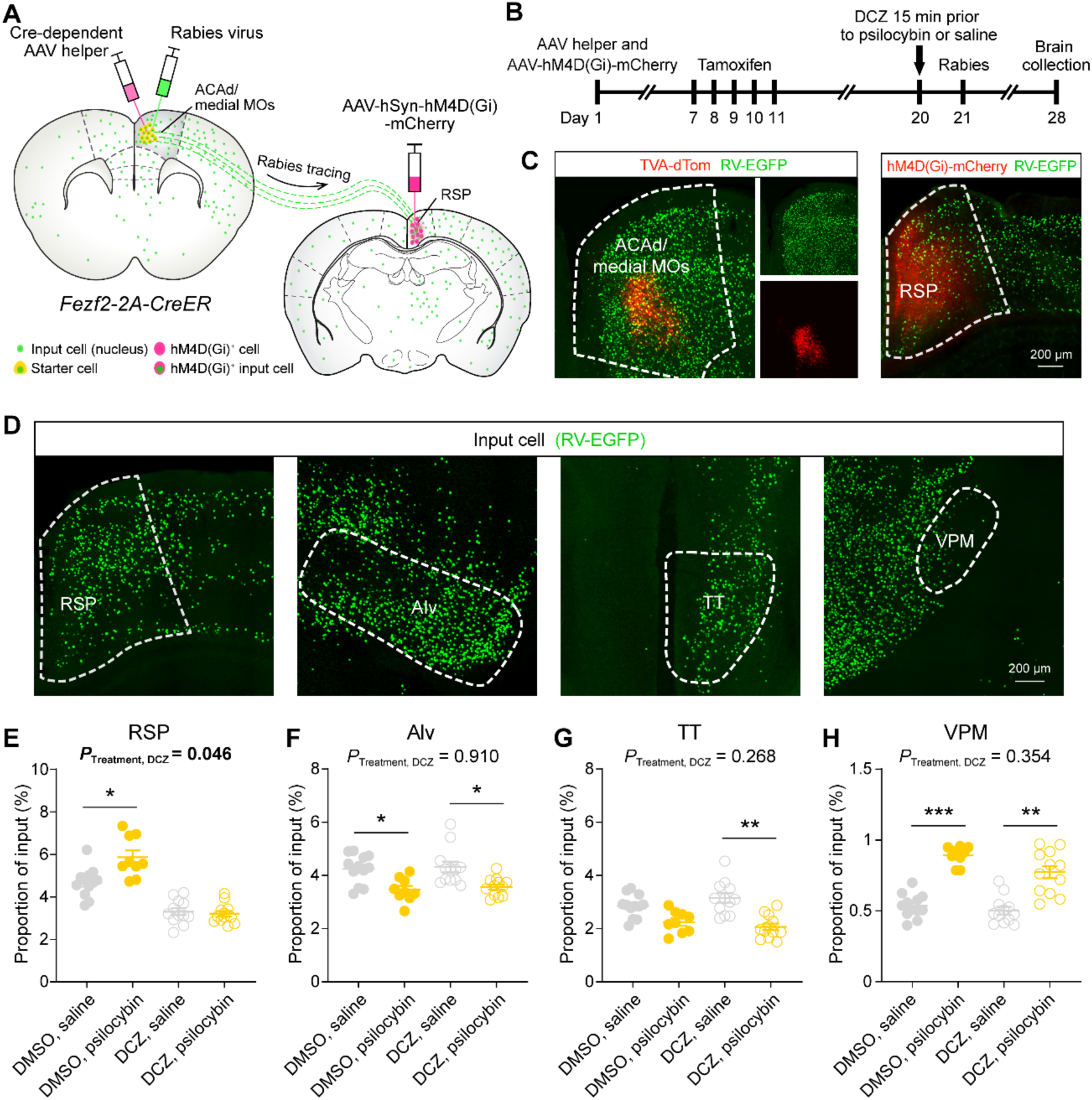
Chemogenetic silencing of retrosplenial cortex alters psilocybin-evoked rewiring of retrosplenial inputs to dorsal medial frontal cortex. **(A)** Experimental setup **(B)** Experimental timeline. **(C)** Fluorescence images from histology after brain collection. TVA-dTom, from Cre-dependent AAV helper virus. RV, rabies virus. **(D)** Fluorescence images showing input cells in RSP, Alv, TT, and VPM. Dashed line, boundary of the brain region. **(E - H)** Effect of DCZ or DMSO and psilocybin or saline on the input cell fraction in RSP, Alv, TT, and VPM. Circle, individual brain slices. Line, mean ± SEM.

We performed the experiment using 3 – 4 animals per condition and obtained 3 brain sections per animal (n = 2 male and 2 female animals for DMSO + saline, 1 male and 2 female animals for DMSO + psilocybin, 2 male and 2 female animals for DCZ + saline, and 2 male and 2 female for DCZ + psilocybin). As anticipated, we observed co-expression of red and green fluorescence in starter cells at the injection site in ACAd and medial MOs (**Figure 6C**). At RSP, we detected green fluorescence in input cells and additionally red fluorescence from the mCherry fluorophore that is fused to the chemogenetic hM4D(Gi) receptor expressed in RSP neurons (**Figure 6C**). For the RSP, there was a significant interaction effect (drug and chemogenetics interaction: *P* = 0.046, linear mixed effects model), indicating that inhibiting neural activity in RSP interfered with the psilocybin-induced potentiation of RSP inputs to the frontal cortical PT^Fezf2^ neurons. In control animals that received DMSO vehicle, psilocybin increased the fraction of input cells (saline: 4.7±0.2%, psilocybin: 5.9±0.3%; *P* = 0.03, post hoc comparison; **Figure 6D, E**). However, this drug-evoked difference in input fraction was abolished when the neural activity in RSP was suppressed via DCZ (saline: 3.3±0.2%, psilocybin: 3.2±0.1%, *P* = 1; **Figure 6D, E**). We note that there was an effect for DCZ relative to control in saline animals (*P* = 0.01; **Figure 6D, E)**, suggesting that lowering the neural activity also affected the efficacy of the monosynaptic tracing independent of the psilocybin administration. Although the coronal sections were intended to target RSP, we could visualize several other presynaptic regions in the same mouse. We determined the input cell fraction in presynaptic regions that previously showed decreases after psilocybin, such as AIv of the lateral network and TT of the ventromedial prefrontal cortex, and found that the RSP manipulation had no influence because psilocybin still weakened these connections (drug and chemogenetics interaction: *P* = 0.9 for AIv and *P* = 0.3 for TT; **Figure 6F, G**). Likewise, the RSP manipulation had no detectable effect on the psilocybin-evoked increase of input fraction in the VPM (drug and chemogenetic interaction: *P* = 0.4; **Figure 6H**). Altogether, the results show that disrupting spiking activity in a presynaptic region influences the psilocybin-evoked rewiring process, specifically for inputs that originate from the manipulated brain region.

## DISCUSSION

We found that psilocybin led to the reorganization of presynaptic inputs into the two major subtypes of pyramidal neurons in the dorsal medial frontal cortex. To conceptualize the overall impact of the changes, it is essential to consider not only the inputs, but also the output targets of the PT and IT neurons. The long-range axonal collaterals of PT and IT neurons in ACAd and medial MOs have recently been mapped at the single-neuron level^54^. PT neurons send axons to numerous destinations outside the cerebrum. IT neurons project to caudoputamen and claustrum, but also to various cortical networks including most prominently to the medial network including RSP and the visual cortex. **Figure 7A** is the synthesis bringing together these output patterns and our data on psilocybin’s effects on presynaptic inputs. The results support two important conclusions. One, psilocybin strengthens the routing of inputs from RSP (a core region in the mouse homolog of default mode network^44–46^), visual cortex, and sensorimotor network to subcortical targets via frontal cortical PT neurons (**Figure 7B**). Interestingly, this shift was done at the expense of reduced influence from the lateral network (including the AIv, hypothesized to be the central seed for the mouse homolog of the salience network^46,55^) and ventromedial prefrontal cortex. Two, psilocybin weakens recurrent loops in the cortex (**Figure 7C**), because inputs from RSP, VIS, and other regions of the medial network are decreased in frontal cortical IT neurons. These are the cortico-cortical targets that IT neurons in the ACAd and medial MOs most strongly innervate, yet their inputs are now reduced after psilocybin.

**Figure 7.**
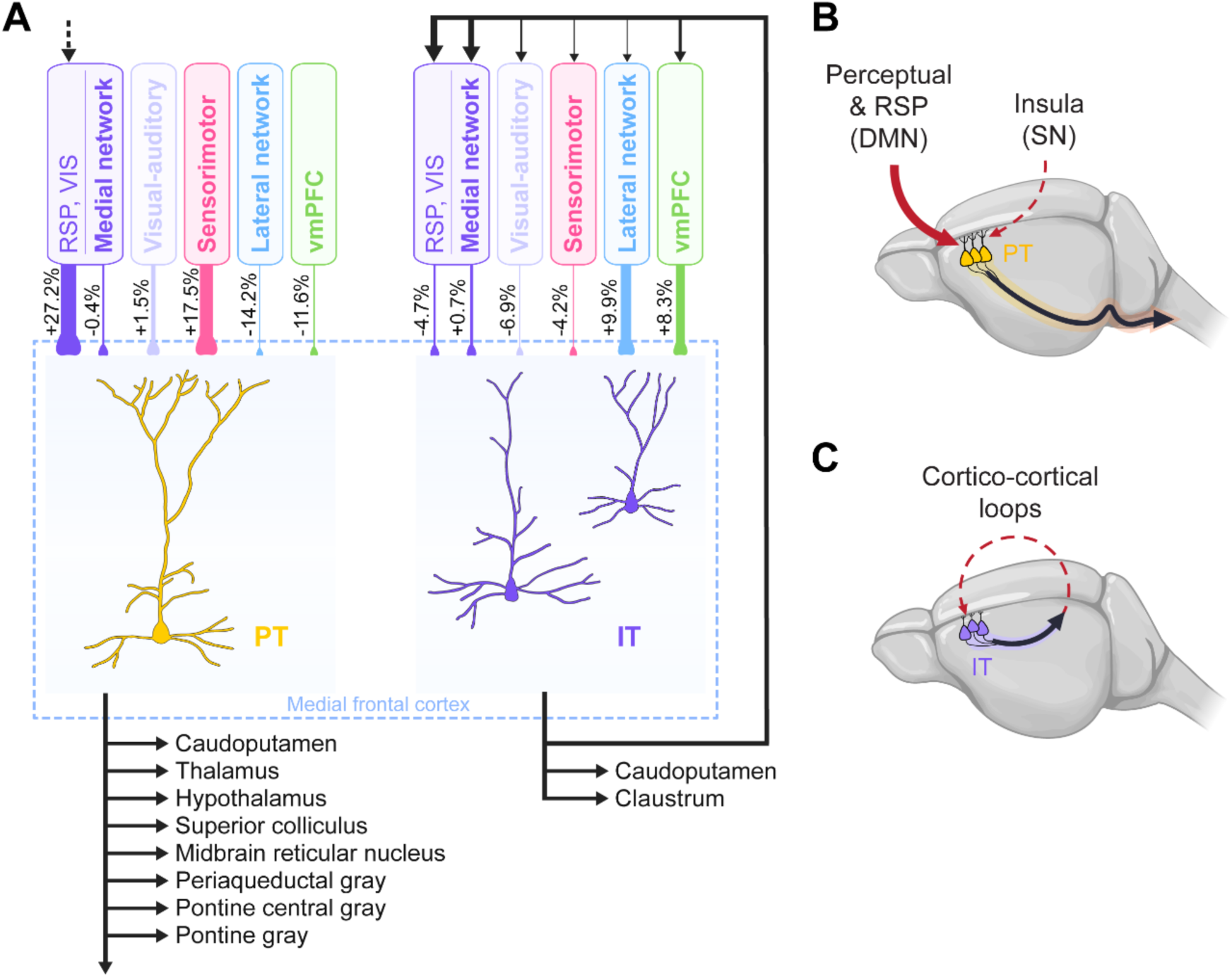
Impact of psilocybin on the connectivity of large-scale cortical networks. **(A)** Schematic depicting the effects of psilocybin on PT^Fezf2^ and IT^PlxnD1^ neurons in the dorsal medial frontal cortex. Inputs to PT^Fezf2^ and IT^PlxnD1^ neurons are drawn with the line thickness directly related to the drug-evoked difference in input fraction based on Figures 2F, 3F, and 3H. Output from IT^PlxnD1^ neurons to the cortical networks are drawn with the line thickness based on Figure 2A of a prior study of single-neuron reconstructions of projection neurons including in the mouse ACAd and medial MOs^54^. Dashed arrow, one subclass of PT neurons in the ACA (although mostly in ACAv) can project to RSP and to a lesser extent VIS^54^. **(B)** Cartoon illustration of the impact on frontal cortical PT^Fezf2^ neurons: strengthen routing of signals from perceptual and retrosplenial regions, mouse homolog of the default mode network, to subcortical target, at the expense of inputs from the insular cortex. **(C)** Cartoon illustration of the impact on frontal cortical IT^PlxnD1^ neurons: weaken connections that are part of the cortico-cortical recurrent loops.

The cell type-specific recording and chemogenetic experiments suggest that spiking activity is a factor that determines if inputs from a presynaptic region would be strengthened by psilocybin. Although the idea may be novel for psychedelic drug action, the results align with a long history of studies demonstrating a crucial role for presynaptic activity in driving the formation of excitatory synapses during neurodevelopment. For example, deprivation of visual stimuli by raising mice in darkness reduces the number of apical dendritic spines in layer 5 neurons in the mouse visual cortex^56^. This deficit is due to the lack of presynaptic activity because the reduction of synapses can also be achieved by blocking action potentials in the optic nerve^57^. By contrast, repeated presentations of visual stimuli can induce long-term potentiation of retinotectal synapses, although presynaptic activity alone was insufficient and the plasticity also requires the firing of postsynaptic neurons^58^. In a similar vein, we surmise that while presynaptic input activity is essential for psilocybin-evoked rewiring, it is likely not sufficient on its own. Instead, we propose that more than one condition may need to be met to strengthen an input: increased presynaptic firing (demonstrated in this study) and elevated dendritic excitability in the postsynaptic neuron^41^, potentially driven by agonism of 5-HT_2A_ receptors^59,60^. For frontal cortical PT neurons, both conditions are present because the cell type exhibits elevated calcium transients in dendritic branches and spines after psilocybin administration, indicating increased dendritic excitability^30^. By contrast, psilocybin does not appear to alter dendritic excitability in frontal cortical IT neurons^30^, leaving the mechanisms underlying input modifications in these cells unclear.

This study demonstrates a potential method to sculpt the psilocybin-evoked synaptic rewiring by manipulating the activity of a presynaptic region. We tested this approach in RSP, which is a central node for the mouse homolog of the default mode network^44–46^. A function of the RSP is to represent perspectives across different spatial reference frames^61^. Psilocybin weakens spatial encoding in RSP neurons in mice^62^, therefore drug action in this retrosplenial cortical region (which is often labeled as the posteromedial cortex and precuneus^63^) may relate to the acute feeling of a temporary loss of one self after psychedelic use in humans. This framework echoes studies pinpointing the deep posteromedial region in mediating the dissociative effects of ketamine^50,51^. Therefore, the long-range connectivity between RSP and dorsal medial frontal cortex is of crucial importance, plausibly serving as a bridge that links the acute subjective changes with the enduring therapeutic effects. In humans, neural activity may be modulated using methods such as repetitive transcranial magnetic stimulation. Our study hints at an exciting avenue for future research to combine neuromodulation with psychedelics to precisely target specific circuits for neural plasticity.

### Limitations of the study

There are limitations in this study. Based on the number of input cells, it is likely that rabies tracing does not label all cells that send monosynaptic inputs to a starter cell. Moreover, it is unknown if rabies virus may favor crossing some synapses over others, for example biases for strong synapses or specific presynaptic cell types. In a prior study, increased labeling of input cells was found in a presynaptic region that has abnormally high neural activity^64^. This should not be a confound in our study, because psilocybin has a short half-life such that the drug-evoked changes in spiking activity would have subsided when the rabies virus was injected on the next day. Due in part to these considerations, it has been argued that rabies tracing should be used as an initial screen to identify putative sources of presynaptic inputs, which then must be confirmed using other methods^65^, which echoes the approach in this study. Furthermore, we used transgenic mice to target PT^Fezf2^ and IT^PlxnD1^ neurons, which capture only a subset of the larger PT and IT cell populations in the medial frontal cortex. Another limitation is that we only tested psilocybin. Synaptic rewiring seems to be a property shared by compounds with putative fast-acting antidepressant effects, because ketamine, 5-MeO-DMT, and other psychedelic-derived novel chemical entities similarly promote dendritic spine formation in the mouse neocortex^13,14,16,66–68^. It remains to be determined how other pharmacological interventions may exert similar or different effects on brain-wide connectivity.

## ACKNOWLEDGEMENTS

We thank Catherine Chong for help with image analysis and Olesia Bilash for advice on setting up photostimulation for slice electrophysiology. Psilocybin was provided by Usona Institute’s Investigational Drug & Material Supply Program; the Usona Institute IDMSP is supported by Alexander Sherwood, Robert Kargbo, and Kristi Kaylo in Madison, WI. The authors acknowledge the support of NIH/NIMH grants R01MH121848, R01MH128217, R01MH137047 (A.C.K.) One Mind–COMPASS Rising Star Award (A.C.K.), NIH training grants T32GM007205 (P.A.D. and N.K.S.), NIH fellowships F30DA059437 (P.A.D.) and F30MH129085 (N.K.S.), and NIH instrumentation grant S10OD032251 (Cornell Biotechnology Resource Center Imaging Facility).

## AUTHOR CONTRIBUTIONS

Q.J. and A.C.K planned the study. Q.J. performed the rabies viral tracing, Neuropixels, and chemogenetics experiments. L.S. performed the two-photon imaging and slice electrophysiology experiments. P.A.D. assisted with recording for the Neuropixels experiment. J.D.N. assisted with histology for the Neuropixels experiment. S.Y. and H.Z. provided the rabies viruses. G.T., T.S.H., H.-M.L., and K.T.B. provided additional reagents. Q.J. analyzed the Neuropixels and chemogenetics data. L.S. analyzed the slice electrophysiology data. N.K.S. performed the transcript and c-Fos analyses. A.D.G. analyzed the two-photon imaging data. A.C.K. analyzed the rabies viral tracing and afferent data. Q.J. and A.C.K. drafted the manuscript. All authors reviewed the manuscript before submission.

## SUPPLEMENTAL TABLES

**Table S1:** Statistical analyses using linear mixed effects models.

## SUPPLEMENTAL FIGURES

**Figure S1.** Control experiments validating monosynaptic tracing in *Fezf2-2A-CreER* mice, related to Figure 1.

**Figure S2.** Estimates for the proportion of excitatory cells as PT, IT, PT^Fezf2^, and IT^PlxnD1^ neurons in the mouse dorsal medial frontal cortex, related to Figure 1, 2.

**Figure S3.** The distribution of input and starter cells in PT^Fezf2^ and IT^Plxnd1^ mice after saline or psilocybin treatment, related to Figure 1, 2.

**Figure S4.** Drug-evoked difference and network selectivity for IT neurons in *PlexinD1-2A-CreER* mice, related to Figure 2.

**Figure S5.** Analysis of the laminar profile of axonal density from presynaptic regions, related to Figure 3.

**Figure S6.** Additional analyses and data for the slice electrophysiology experiments, related to Figure 4.

**Figure S7.** Additional analyses for the Neuropixels experiments, related to Figure 5.

## SUPPLEMENTAL VIDEOS

**Video S1:** Light sheet fluorescence images showing the brain-wide distribution of input cells (green) into the frontal cortical PT^Fezf2^ starter cells (red and green) in a *Fezf2-2A-CreER* mouse after psilocybin administration, related to Figure 2.

**Video S2:** Light sheet fluorescence images showing the frontal cortical PT^Fezf2^ neurons (red) and NeuN stain (white) in a *Fezf2-2A-CreER* mouse after psilocybin administration, related to Figure 2.

**Video S3:** Light sheet fluorescence images showing the brain-wide distribution of input cells (green) into the frontal cortical IT^PlxnD1^ starter cells (red and green) in a *PlexnD1-2A-CreER* mouse after psilocybin administration, related to Figure 3.

**Video S4:** Light sheet fluorescence images showing the frontal cortical IT^PlxnD1^ neurons (red) and NeuN stain (white) in a *PlexnD1-2A-CreER* mouse after psilocybin administration, related to Figure 3.

## RESOURCE AVAILABILITY

### Lead contact

Further information and requests for resources and reagents should be directed to and will be fulfilled by the Lead Contact Alex C. Kwan.

### Materials availability

This study did not generate new unique reagents.

### Data and code availability

The data that support the findings and the code used to analyze the data in this study will be made publicly available at https://github.com/Kwan-Lab.

## EXPERIMENTAL MODEL AND STUDY PARTICIPANT DETAILS

### Animals

*Fezf2-2A-CreER* (B6;129S4-*Fezf2^tm1.1(cre/ERT2)Zjh^*/J, Stock No. 036296)^25^, *PlexinD1-2A-CreER* (B6;129S4-*Plxnd1^tm1.1(cre/ERT2)Zjh^*/J, Stock No. 036294)^25^, *Ai32* (B6.Cg-*Gt(ROSA)26Sor^tm32(CAG-COP4*H134R/EYFP)Hze^*/J, Stock No. 024109), *Ai14* (B6.Cg-*Gt(ROSA)26Sor^tm14(CAG-tdTomato)Hze^*/J, Stock No. 007914), *Ai75* (B6.Cg-*Gt(ROSA)26Sor^tm75.1(CAG-tdTomato*)Hze^*/J, Stock No. 025106) and wild-type C57BL/6J (Stock No. 000664) mice were acquired from Jackson Laboratory and bred in our animal facility. For tracing experiments involving G-deleted rabies virus and adeno-associated virus (AAV), homozygous 2- to 3-month-old *Fezf2-2A-CreER* and *PlexinD1-2A-CreER* mice of both sexes were used. For Neuropixels studies, 2- to 3-month-old *Ai32* mice of both sexes were used. For two-photon imaging studies, 5- to 7-week-old mice of both sexes were used. Animals were housed in groups of 2 to 5 per cage in a temperature-controlled room, maintained on a standard 12-hour light/dark cycle (lights on from 8:00 AM to 8:00 PM). Food and water were provided *ad libitum*. Animals were randomly assigned to experimental groups. All animal care and experimental procedures were approved by the Institutional Animal Care and Use Committee (IACUC) at Cornell University.

## METHOD DETAILS

### Animal numbers

For monosynaptic tracing, we had 9 *Fezf2-2A-CreER* animals for psilocybin condition (4 male and 5 female), 8 *Fezf2-2A-CreER* animals for saline condition (4 male, 4 female), 8 *PlexinD1-2A-CreER* animals for psilocybin condition (4 male, 4 female), 8 *PlexinD1-2A-CreER* animals for saline condition (4 male, 4 female). For in vivo electrophysiology, we used 6 *Ai32* animals for the psilocybin condition (2 male, 4 female) and 6 *Ai32* animals for the saline condition (3 male, 3 female). For chemogenetics, we used 4 animals in the DMSO + saline group (2 male, 2 female), 3 animals in the DMSO + psilocybin group (1 male, 2 female), 4 animals in the DCZ + saline group (2 male, 2 female), and 4 animals in the DCZ + psilocybin group (2 male, 2 female). For two-photon imaging, we used 7 *Fezf2-2A-CreER* animals for the saline condition (2 male, 5 female) and 8 *Fezf2-2A-CreER* animals for the psilocybin condition (4 male, 4 female), with a total of 60 and 62 fields of view imaged for saline and psilocybin conditions, respectively. For immunohistochemistry, we used 4 *Fezf2-2A-CreER;Ai75* animals (4 females) and 5 *PlexinD1-2A-CreER;Ai75* animals (5 females). For slice electrophysiology, we used two strategies to label the PT^Fezf2^ neurons. For transgenic Ai14-based labeling, we used 6 *Fezf2-2A-CreER;Ai14* animals (2 male, 4 female) for the saline condition at 24 hr time point, 7 *Fezf2-2A-CreER;Ai14* animals (4 male, 3 female) for the psilocybin condition at 24 hr time point, 6 *Fezf2-2A-CreER;Ai14* animals (3 male, 3 female) for the saline condition at 3 day time point, and 7 *Fezf2-2A-CreER;Ai14* animals (5 male, 2 female) for the psilocybin condition at 3 day time point. For viral mediated tdTomato expression, we used 4 *Fezf2-2A-CreER* animals (1 male, 3 female) for the saline condition and 5 *Fezf2-2A-CreER* animals (2 male, 3 female) for the psilocybin condition.

### Viruses

For rabies viral tracing, the Cre-dependent AAV helper (AAV1-hSyn-DIO-TVA^66T^-dTomato-CVS N2c G), Cre-dependent AAV control (AAV1-hSyn-DIO-TVA^66T^-dTomato), and EnvA-pseudotyped G-deleted rabies virus (EnvA-CVS N2c^ΔG^-H2B-EGFP) were generated at the Allen Institute as previously described^22^. For in vivo electrophysiology, AAVretro-hSyn-Cre-WPRE-hGH (Catalog #105553) was purchased from Addgene. For chemogenetics, AAV9-hSyn-hM4D(Gi)-mCherry (Catalog #50475) was purchased from Addgene. For two-photon imaging, AAV1-pCAG-FLEX-EGFP-WPRE (Catalog #51502) was purchased from Addgene, and AAV9-hSyn-Synaptophysin-mRuby2-WPRE-polyA vector was custom-designed and packaged by a commercial vendor (BrainVTA). For immunohistochemistry, AAV9-mDlx-GFP-Fishell-1 (Catalog #83900) was purchased from Addgene. For slice electrophysiology, AAV1-FLEX-tdTomato (Catalog #28306) and AAV9-hSyn-hChR2(H134R)-EYFP (Catalog #26973) were purchased from Addgene. The viruses were aliquoted and stored at -80°C. Prior to stereotaxic injection, the viruses were removed from the -80°C freezer and thawed on ice.

### Surgery

Surgery commenced with anesthesia induction using 2-3% isoflurane. Once the animal reached a surgical level of anesthesia, it was placed in a stereotaxic frame (Model 900, David Kopf Instruments). Dexamethasone (2 mg/kg, i.m.; #1DEX022, Bimeda) and carprofen (5 mg/kg, s.c.; #059149, Covetrus) were administered for anti-inflammatory and analgesic effects. Eye lubricant (Optixcare Eye Lube, #062143, Covetrus North America) was applied to protect the eyes from drying and potential damage. Anesthesia was maintained at 1-1.5% isoflurane during the entire procedure. Core body temperature was maintained at 38°C using a far-infrared warming pad (#RT-0515, Kent Scientific). The area above the skull was shaved. The scalp was sterilized with ethanol pads and povidone-iodine. A small incision (∼1 cm) was made along the midline. The skin and fascia were carefully removed to expose the skull. Burr holes were drilled at specific target sites using a handheld dental drill (#HP4-917, Foredom). The viruses were injected intracranially via a borosilicate glass micropipette connected to an injection control unit (Nanoject II Auto-Nanoliter Injector, Drummond Scientific). Injections were performed for various experiments using different viral vectors and volumes, as detailed below, using 4.6 nL pulses with a 20-second interval between pulses. To minimize backflow, the micropipette was left in place for 5-10 minutes following the end of the injection before being withdrawn slowly. The brain surface was kept moist with artificial cerebrospinal fluid (aCSF; in mM: 135 NaCl (#S5150, Sigma-Aldrich), 5 HEPES (#H3537, Sigma-Aldrich), 5 KCl (#60142, Sigma-Aldrich), 1.8 CaCl_2_ (#21115, Sigma-Aldrich), 1 MgCl_2_ (#63069, Sigma-Aldrich); pH 7.3). After completing the injections, the burr holes were covered with a silicone elastomer (#10006546, Smooth-On, Inc.). The incision was sealed using surgical sutures (#1265B, Surgical Specialties Corporation). Post-surgical analgesia was provided with carprofen (5 mg/kg, subcutaneously) immediately following the procedure and once daily for three days thereafter.

All mice received injections in one hemisphere only, and we targeted the right hemisphere. The anteroposterior (AP) and mediolateral (ML) coordinates are measured from bregma, with ML distances taken from the midline. The depth was measured from the brain’s pial surface. For monosynaptic tracing of inputs, on day 1, the Cre-dependent AAV helper (276 nL) was injected into the medial frontal cortex (coordinates are AP: 1.6 mm, ML: 0.4 mm, depth: -0.6 and -0.8 mm) in a *Fezf2-2A-CreER* or *PlexinD1-2A-CreER* mouse. More specifically, half of the volume was injected at a depth of -0.6 mm and the remainder was injected at -0.8 mm. The anatomical coordinates correspond to the ACAd (anterior cingulate area, dorsal part) and the medial portion of MOs (secondary motor area) of the mouse. For control experiments, the Cre-dependent AAV control (276 nL) was injected instead. On days 7 through 11, tamoxifen was administered daily to induce the expression of Cre recombinase. On day 20, psilocybin (1 mg/kg, i.p.; prepared fresh monthly from powder; Usona Institute) or saline was administered. On day 21, G-deleted rabies virus (276 nL) was injected in the medial frontal cortex at the same stereotaxic coordinates. On day 28, the mouse was deeply anesthetized for transcardial perfusion.

For in vivo electrophysiology, the AAVretro-hSyn-Cre-WPRE-hGH (276 nL) was injected into the medial frontal cortex (AP: 1.6 mm, ML: 0.4 mm, depth: -0.6 and -0.8 mm) of an *Ai32* mouse. Three weeks later, the mice underwent a second surgical procedure. A midline incision was made, the skin above the skull was removed, and the periosteum of the skull was cleared. A 0.9-mm-diameter craniotomy was drilled using a handheld dental drill, and a self-tapping bone screw (0.86 mm; #19010-10, Fine Science Tools) was placed into the cerebellum to serve as a ground screw. A stainless steel headplate (eMachineShop; design available at https://github.com/Kwan-Lab/behavioral-rigs) was affixed to the skull using a rapid adhesive cement system (C&B Metabond, #S380, Parkell). The mouse would recover for at least one week before commencing the electrophysiological recording.

For chemogenetics, on day 1, the Cre-dependent AAV helper virus (diluted 1:3 in phosphate-buffered saline (PBS; #P4474, Sigma-Aldrich); 276 nL) was injected into the medial frontal cortex (AP: 1.6 mm, ML: 0.4 mm, depth: -0.6 and -0.8 mm) and AAV9-hSyn-hM4D(Gi)-mCherry (276 nL) was injected into the retrosplenial cortex (RSPd, retrosplenial area, dorsal part, and RSPv, retrosplenial area, ventral part; AP: -1.7 mm, ML: 0.15 mm, depth: -0.6 and -0.8 mm) in a *Fezf2-2A-CreER* mouse. On days 7 to 11, tamoxifen was administered daily to induce Cre-dependent gene expression. On day 20, deschloroclozapine (DCZ; 0.1 mg/kg, i.p.; #HY-42110, MedChemExpress) was administered from a stock solution (5 mg/mL) prepared in 100% DMSO and diluted with saline to achieve a final injection concentration (0.025 mg/mL) containing 0.5% v/v DMSO. For control, an equivalent amount of DMSO (0.5% v/v in saline) was administered. Then 15 minutes later, psilocybin (1 mg/kg, i.p.) or saline (i.p.) was injected. On day 21, the G-deleted rabies virus (276 nL) was injected in the medial frontal cortex at the same stereotaxic coordinates. On day 28, the mouse was deeply anesthetized and perfused transcardially with PBS followed by 4% paraformaldehyde (4% (v/v) in PBS). The brain was extracted and post-fixed in 4% paraformaldehyde overnight at 4°C.

For two-photon imaging of axonal boutons and dendritic structures, viral injections and window implantation were performed on the same day. Following induction of anesthesia, a midline scalp incision was made, and the skin above the skull was removed. The skull surface was gently cleared of connective tissue. AAV9-hSyn-Synaptophysin-mRuby2-WPREpolyA (276 nL) was injected into the retrosplenial cortex (AP: -1.7 mm, ML: 0.15 mm, depth: -0.6 and -0.8 mm) of a *Fezf2-2A-CreER* mouse. Next, a 3 mm circular craniotomy was created over the medial frontal cortex (centered at AP: 1.6 mm, ML: 0.4 mm) using a dental drill. The exposed dura was kept moist with aCSF throughout the procedure. AAV1-pCAG-FLEX-EGFP-WPRE (1:100 dilution in PBS; 92 nL) was then injected into the medial frontal cortex (AP: 1.6 mm, ML: 0.4 mm, depth: -0.6 and -0.8 mm). Following viral injection, a double-layer glass window was placed over the craniotomy. The window was constructed by bonding two round coverslips (3 mm diameter, 0.15 mm thickness; #64-0720, Warner Instruments) using ultraviolet-curable optical adhesive (#NOA 61, Norland Products), which was cured with an ultraviolet illuminator (#2182210, Loctite). Super glue adhesive (Henkel Loctite 454) was applied with caution to secure the window to the surrounding skull while maintaining slight pressure. A stainless steel headplate was then fixed onto the skull and centered on the glass window using a rapid adhesive cement system (C&B Metabond, Parkell). The mouse would recover and ensure sufficient AAV expression for at least 3 weeks after the window implantation and AAV injection before beginning imaging experiments.

To achieve CreER-induced gene expression, tamoxifen (#T5648, Sigma-Aldrich) was prepared as a 20 mg/mL solution in corn oil (#C8267, Sigma-Aldrich). The solution was fully dissolved using an ultrasonic bath at 37°C for 1-2 hours, then aliquoted into 1 mL portions, protected from light with aluminum foil, and stored at -20°C. Before administration, the tamoxifen aliquots were thawed and kept at 4°C. Each mouse was weighed, and tamoxifen was injected intraperitoneally at a dose of 75 mg/kg (equivalent to 0.094 mL of the 20 mg/mL solution for a 25 g mouse) once daily for five consecutive days (https://www.jax.org/research-and-faculty/resources/cre-repository/tamoxifen). The injection site was sanitized with 70% ethanol before each injection.

### Whole-brain imaging after monosynaptic tracing: Light sheet fluorescence microscopy

Each mouse was perfused transcardially with ice-cold PBS containing 10 U/mL heparin (H3149, Sigma-Aldrich) and ice-cold 4% paraformaldehyde (PFA). After perfusion, the brain was extracted and incubated in 4% PFA at 4°C for 24 hours, then washed twice with PBS. The samples were stored in PBS with 0.02% sodium azide (#BDH7465-2, VWR) and then shipped to LifeCanvas Technologies (Cambridge, MA) for tissue clearing and imaging. The brains were preserved with SHIELD and then cleared with SmartClear technology, then immunolabeled for NeuN (#MCA-1B7, Encor Biotechnology) with eFLASH technology, which combined stochastic electrotransport and SWITCH techniques within a SmartBatch device. Following labeling, samples were incubated overnight at 37°C in 50% EasyIndex (RI = 1.52, LifeCanvas Technologies), then further incubated for 24 hours in 100% EasyIndex to achieve complete refractive index matching. For consistent index matching, the samples were embedded in 2% ultra-low melt agarose prepared with EasyIndex and left in EasyIndex overnight. Imaging was performed on a SmartSPIM axially-swept light sheet microscope with a 3.6x objective (0.2 NA), using a z-step size of 4 μm and an XY pixel size of 1.8 μm. The samples are imaged using three channels: 488 nm (GFP), 561nm (dTomato), and 647 nm (NeuN).

### Whole-brain imaging after monosynaptic tracing: Image processing and cell detection

Each sample was aligned to the Allen Brain Atlas (Allen Institute: https://portal.brain-map.org/) through an automated alignment process conducted by LifeCanvas Technologies. NeuN channels for individual brains were registered to an average NeuN atlas generated by LifeCanvas Technologies using previously aligned samples. The registration was performed in stages using rigid, affine, and b-spline transformations (SimpleElastix: https://simpleelastix.github.io/). Cell detection was carried out through a two-step neural network approach. Initially, a U-Net-based fully convolutional network^69^ was employed to detect potential cell locations with high sensitivity. Following this, a second network based on a ResNet architecture^70^ was used to classify each detected location as either positive or negative, refining the accuracy of cell detection. This model leveraged both U-Net and Fully Convolutional Network architectures to optimize semantic segmentation. For colocalization analysis, single-channel cell centers were identified, and the spatial relationships between cells across channels were calculated. Cells in different channels were marked as colocalized if the Euclidean distance between a cell in channel 1 and a cell in channel 2 was less than three voxels, indicating coexpression within a single cell. Data after imaging processing and cell detection were tabulated in CSV files, which included region IDs and cell counts in each region for *Fezf2-2A-CreER* and *PlexinD1-2A-CreER* mice. We divided the brain into the 316 non-overlapping regions based on the “summary structures” defined by a prior study of the Allen Common Coordinate Framework (CCF)^31^.

### Whole-brain imaging after monosynaptic tracing: Analysis of starter cells

Starter cells were defined as those nuclei detected with coexpression in the 488 and 561 nm channels and located in one of the frontal cortical regions in the right hemisphere (MOp-R, ACAv-R, MOs-R, ACAd-R, PL-R, ILA-R, ORBm-R, and ORBvl-R).

As a quality check, we analyzed cells with colocalized red and green fluorescence in the entire brain (henceforth refer to as “red+green cells”), including in the right frontal cortex as well as also all other brain regions. For each region in each brain, we calculated the fraction of red+green cells, by dividing the number of red+green cells in a region by the total number of red+green cells in the brain. Unexpectedly, we observed a sizable number of puncta with red and green fluorescence in the left and right main olfactory bulb (MOB). We investigated this further by looking at the raw fluorescence images, which showed that the red-green puncta were in the glomerular layer of the olfactory bulb. These red-green puncta did not colocalize with NeuN, indicating that the signals did not originate from neurons, and instead likely come from lipofuscin granules^71^. When we excluded these red+green puncta in the MOB from our analysis, we found that nearly all (91.1%) of the red+green cells were indeed found in the right frontal cortex (MOp-R, MOs-R, ACAd-R, ACAv-R, PL-R, ILA-R, ORBl-R, ORBm-R, and ORBvl-R). Outside the right frontal cortex, the top three highest contributors were neighboring regions in the right anterior cortex with 1.01% of the red+green cells coming from FRP-R, 0.49% coming from DP-R, and 0.43% coming from ENTI-R. There was negligible signal in the contralateral hemisphere, because only 0.54% of the red-green cells were detected in left frontal cortex (MOp-L, MOs-L, ACAd-L, ACAv-L, PL-L, ILA-L, ORBl-L, ORBm-L, and ORBvl-L). Overall, the starter cells were localized to the injection site as expected.

### Whole-brain imaging after monosynaptic tracing: Analysis of input cells

Inputs cells were defined as those nuclei detected in the 488 nm. For the frontal cortical regions in the right hemisphere, some nuclei detected in 488 nm were also the nuclei with coexpression in the 488 and 561 nm channels, so the count was subtracted by the number of starter cells to yield the number of input cells.

The number of input cells in a brain varied depending on the number of starter cells, and the relationship was not linear. To aggregate the results across brain, we normalized by calculating, for each region of each brain, the “proportion of total input (%)” by dividing the number of input cells in the region by the total number of input cells in the brain. Next, instead of keeping track of all the summary structures, we focused on the regions that provided substantial inputs to the medial frontal cortex. We set a threshold to identify regions that contributed at least 0.3% of the total input cells for any experimental group (PT or IT, psilocybin or saline). This yielded regions that provided substantial inputs, which we called “presynaptic regions”.

To evaluate the effect of psilocybin, for each presynaptic region, we computed “drug-evoked difference (%)” by subtracting the mean input fraction for psilocybin by the mean input fraction for saline, and normalized by divided by the mean input fraction for saline. To estimate the variance in drug-evoked difference, for each presynaptic region, we performed bootstrapping by drawing with replacement 75% of the psilocybin samples and 75% of saline samples, then calculated the drug-evoked difference for this scenario for 10,000 repeats. Using the distribution of drug-evoked differences from the bootstrapping, we estimate the 90% confidence interval for the drug-evoked difference for a presynaptic region.

To assess network-specific effects, we grouped presynaptic regions based on their network memberships. Networks were defined based on a prior large-scale connectivity study of the mouse neocortex^18^, including 5 networks: medial (ACAd, ACAv, RSPagl, RSPd, RSPv, VISa, VISrl), visual-auditory (AUDd, AUDp, AUDpo, AUDv, VISal, VISam, VISl, VISp, VISpl, VISpm, VISli, VISpor), sensorimotor (MOp, MOs, SSp-n, SSp-bfd, SSp-ll, SSp-m, SSp-ul, SSp-tr, SSp-un, SSs), lateral (GU, VISC, AId, AIp, AIv, TEa, PERI, ETC, PIR, ENTI), and ventromedial prefrontal cortex (PL, ILA, ORBl, ORBm, ORBvl, AON, TT, DP). Some presynaptic regions showed increased number of input cells after psilocybin relative to saline, whereas other presynaptic regions showed decreased number of input cells. To determine if the distribution of regions with increase or decrease (2 categories) are biased or evenly distributed for the networks (5 categories), we used the chi-squared test.

To determine cell-type differences in the network-specific effects, presynaptic regions that do not belong to these networks are considered ‘Outside neocortex’. When assessing only saline data to determine baseline differences for inputs into PT and IT neurons, we would sum the number of input cells in the presynaptic regions that belong to a network. We used the two-sided Wilcoxon rank sum test to determine statistical significance between the two cell types for each network. To evaluate the effect of psilocybin on each network, we would first sum the number of input cells in the presynaptic regions that belong to a network, then compute “drug-evoked difference (%)” by subtracting the mean input fraction for psilocybin by the mean input fraction for saline, and normalized by divided by the mean input fraction for saline. To estimate the variance in drug-evoked difference, for each network, we performed bootstrapping by drawing with replacement 75% of the psilocybin samples and 75% of saline samples, then calculated the drug-evoked difference for this scenario for 10,000 repeats. Using the distribution of drug-evoked differences from the bootstrapping, we estimate the 90% confidence interval for the drug-evoked difference for a network. To determine cell-type specific differences, for each of the 10,000 repeats, we also calculated the difference in the drug-evoked difference between PT and IT neurons. Using this distribution of PT minus IT drug-evoked difference, we deemed the cell-type difference to be significant if 0 lies outside the 95% confidence interval.

For the network selectivity analysis, take PT^Fezf2^ neurons as an example, we started with the 65 presynaptic regions, each of which was associated with a drug-evoked difference. We analyzed the subset of 47 presynaptic regions that are part of the neocortex, with each belonging to 1 of 5 possible networks (medial, visual-auditory, sensorimotor, lateral, vmPFC). Based on the drug-evoked difference, we classified each of the 47 presynaptic regions as an increase or a decrease. A chi-squared test was used to determine if the observing proportions deviate from the expected proportions (e.g., observing a few people accumulating all the wealth while expecting wealth distributed equally). To calculate the chi-square statistic, we needed to know the observed proportions and expected proportions. The observed 5 proportions were the number of regions with increase divided by the number of regions in the 5 networks. The expected 5 proportions were an equal distribution with regions with increase divided evenly among the 5 networks. We determined the chi-square statistic for our experimental data. Based on the chi-square statistic and the degree of freedom ((2-1)*(5-1) = 4), we calculated the associated p-value. To evaluate the null distribution of chi-square statistics, we used bootstrapping. We created surrogate data by randomly shuffling the increase and decrease labels for the 47 presynaptic regions. Then we calculated the chi-square statistic using the shuffled data. We repeated this procedure for 1,000,000 times to generate the histogram corresponding to the null distribution.

### Axonal projection analysis

The Allen Mouse Brain Connectivity Atlas has a curated database of fluorescence images of coronal sections of a mouse brain after injecting viral tracer in various brain regions^38^. For each of the 65 presynaptic regions, we identified an experiment in the database in which the injection was done centering at that region. Then we downloaded the fluorescence image corresponding to the medial frontal cortex. Using the ImageJ software, we would draw one rectangle region of interest for ACAd, roughly corresponding to a segment spanning all the cortical layers in ACAd, and extract the pixel intensity along that laminar profile. We would draw another rectangle region of interest for medial MOs and repeat to extract the pixel intensity along that laminar profile.

For analysis, we first interpolate so the length of the ACAd profile matches the length of the MOs profile, because ACAd is thinner than medial MOs. We then average the ACAd and MOs profiles to get a single pixel intensity profile across cortical depth. Based on the pixel values as a function of depth, we fitted a cubic smoothing spline to estimate the axonal density as a function of depth. We defined the laminar positions as follows: layer 1 (0 – 5% of full depth), layer 2/3 (5 – 25%), layer 5 (25 – 50%), layer 6a (50 – 90%), and layer 6b (90 – 100%). Pearson correlation coefficients were used to assess the association between aspects of the axonal density and the drug-evoked difference in PT neurons as measured via monosynaptic tracing.

### Serotonin receptor distribution and c-Fos analyses

Drug-related differences in the rabies tracing data were compared to publicly available region-level data on 5-HT receptor expression patterns and immediate-early gene (c-Fos) activation following psilocybin. For 5-HT receptor expression, *in situ* hybridization (ISH) data for *Htr1a*, *Htr2a*, and *Htr2c* mRNA were extracted across the cortical regions in prior studies^27,40,41^, with the original data coming from the Allen Mouse Brain Atlas gene expression database^39^. Per transcript, expression levels were computed as the mean ISH intensity across voxels of a given cortical region (“ISH energy”), then z-scored across regions for normalization^40^. For c-Fos expression, we used prior data from whole-brain light sheet imaging of c-Fos immunohistochemistry after treatment with psilocybin (1 mg/kg) or saline^42^. For each cortical brain region, we computed “drug-evoked difference (%)” by subtracting the mean c-Fos expression for psilocybin by the mean c-Fos expression for saline, and normalized by dividing the mean c-Fos expression for saline. Regional drug-evoked differences in the rabies tracing data were then compared to these independent estimates of 5-HT receptor expression and drug-evoked differences in c-Fos.

### In vivo electrophysiology: Recording procedure

Mice were acclimated to head fixation over a four-day period, with exposure times progressively increased from 15 minutes on the first day to 30, 60, and finally 120 minutes on the subsequent days. On the day of the experiment, 3 hours before the recording, the mouse was anesthetized with isoflurane and a 2-mm-diameter craniotomy was performed over the retrosplenial cortex (RSP; AP: -1.7 mm, ML: 0.15 mm). To minimize tissue damage and prevent overheating during drilling, cold (4°C) artificial cerebrospinal fluid (aCSF) was applied to clear debris, and care was taken to prevent bleeding and remove bone particles. Following the craniotomy, an area of the dura mater was removed using a fine metal pin (#10130-10, Fine Science Tools). A section of Surgifoam (#1972, Johnson & Johnson), soaked in aCSF, was positioned over the exposed cortex to maintain moisture, and a layer of silicone elastomer (#10006546, Smooth-On, Inc.) was applied to ensure cleanliness and humidity in the craniotomy area until recording commenced. An intravenous catheter system (22-gauge; #383323, BD Saf-T-Intima Closed IV Catheter Systems), preloaded with psilocybin or saline, was kept at neutral pressure and implanted into the intraperitoneal cavity an hour prior to recording. The catheter was secured with Vetbond tissue adhesive (#1469, 3M Vetbond), with the tubing affixed to the head-fixation support frame to ensure stability during the session. Immediately before recording, the silicon polymer and Surgifoam were removed, and the craniotomy was gently rinsed with aCSF to ensure a clear field. A ground wire was wrapped around the self-tapping bone screw placed above the cerebellum. A Neuropixels 1.0 high-density silicon probe (#Neuropixels 1.0, IMEC), configured to record from the bottom-most 384 sites out of 960 available electrodes. Prior the insertion, the probe was coated with a 10 µL droplet of CM-DiI dye (1 mM in ethanol; #C7000, Invitrogen). The probe was inserted at a controlled rate of 100 µm/min via a micromanipulator (MPM; M3-LS-3.4-15-XYZ-MPM-Inverted, New Scale Technologies) until reaching a target depth of approximately 2000 µm. During probe implantation, aCSF was periodically applied to the craniotomy site. Once reaching the final position, the probe was held steady to stabilize for at least 30 minutes before recording commenced. Data were collected using the OpenEphys software in external reference mode, with action potentials recorded between 0.3-10 kHz at 30 kHz and LFP between 0.5-500 Hz at 2.5 kHz. The recording protocol involved a baseline period of 30 minutes, after which mice received an injection of either psilocybin or saline via the catheter, followed by an additional 60 minutes of recording. At the conclusion of each session, optotagging was conducted to identify ChR2-expressing neurons. A 473 nm laser (Obis FP 473LX, Coherent) connected to a 200 µm optical fiber was directed toward the craniotomy site using the micromanipulator, with laser pulses controlled by OpenEphys and triggered via a PulsePal (#1102, Sanworks), delivering 20 ms pulses at 1 Hz with an intensity of ∼25 mW/mm². Each trial lasted 1 second, with a 980 ms inter-trial interval, across a minimum of 500 trials. Following the completion of recording, the probe was carefully retracted and cleaned using a sequence of 1% Tergazyme solution (#1304-1, Alconox Inc.), deionized water, and isopropyl alcohol (#AB07015, AmericanBio).

### In vivo electrophysiology: Data analysis

Raw extracellular traces from the action potential band were processed with SpikeInterface^72^, which provided an end-to-end workflow for filtering, channel clean-up, spike detection, and quality-metric extraction. The preprocessing steps were: (1) passing the traces through a high-pass filter at 400 Hz to remove slow fluctuations while retaining action potentials; (2) excluding channels whose root-mean-square noise exceeded a session-specific threshold, which eliminated mostly the sites that were positioned outside the brain; (3) correcting for phase alignment to compensate for temporal offsets between channels; (4) removing shared noise sources, including specifically the laser-induced artifacts, by common-median referencing. Spike sorting was then performed with Kilosort 2.5^73,74^, which iteratively matched template waveforms and tracked electrode drift, yielding an initial set of putative single units. These putative units were imported into Phy (https://github.com/kwikteam/phy) for manual curation: each cluster was inspected for waveform consistency, refractory-period violations, and stability over time, and subsequently labelled as good (well-isolated single unit), MUA (multi-unit activity), or noise. Only putative units marked good were retained for subsequent analyses. Quality metrics were then generated via SpikeInterface. Units were retained if they satisfied the following criteria^75^: (1) presence ratio ≥ 0.90. Presence ratio refers to the fraction of 60-s time bins in which the unit fired at least one spike, indicating stable activity throughout the session; (2) ISI violation rate < 0.5. ISI violation rate refers to the proportion of spikes occurring within 1.5 ms of a previous spike, indicating refractory-period compliance; (3) amplitude cutoff < 0.1. Amplitude cutoff estimated the fraction of spikes missed because their amplitudes fell below the detection threshold in the sorting algorithm and thus indicated waveform consistency; (4) isolation distance > 20. Isolation distance quantified the separation between the unit’s waveforms and those of other units, ensuring adequate cluster isolation. To identify opto-tagged neurons, peri-stimulus time histograms (PSTHs) were created by aligning the spiking activity of each unit to the onset of laser stimulation. Opto-tagged neurons were then classified based on latency to spike and the reliability of spiking following laser onset.

Raw extracellular traces from the LFP band were analyzed in Python using custom scripts built upon numpy, scipy, and pynapple. Signal from each channel was low-pass filtered (<300 Hz), zero-centered by subtracting the time-averaged mean, and notch filtered to remove potential power-line noise at 60 Hz. LFP signals were segmented into non-overlapping 2-second epochs. Animal’s movements could induce artifacts; therefore, we had a pre-processing step to exclude epochs with outliers. We had two thresholds. One threshold was fixed at ±1800 µV. The other threshold was channel-specific, set to be ±6× the standard deviation as calculated using the channel’s signal. Any epoch that had spurious large-amplitude deflections that lied beyond either of the threshold would be excluded from further analysis. For the remaining epochs, power spectral density was computed using a custom implementation of the multi-taper method (time-bandwidth product = 3.0; number of tapers = 5), based on discrete prolate spheroidal sequences and fast Fourier transforms. Spectral power was calculated for the 1–100 Hz range and averaged across tapers. We made rough estimates of the cortical depth associated with each channel by using Phy to mark the location at which we began to switch from no detected units to detection of well-isolated single units and then calculating depth based on the known spacing between channels. For each range of depths (i.e., 0-300, 300-600, and 600-900 µm), we averaged the spectral power from all channels within that range. We collated the results for pre-drug (-30 – 0 min) and post-drug (0 – 60 min) windows. Subsequently, mean spectral power was extracted from five frequency bands: slow (0.2–1.5 Hz), delta (1.5–4 Hz), theta (4–10 Hz), beta (10–30 Hz), and gamma (30–80 Hz)^76^.

### In vivo electrophysiology: Histology

To evaluate fluorescence at specific brain regions, the fixed brain was sectioned at a thickness of 45 µm using a vibratome (VT1000S, Leica). The coronal sections were mounted on slides with mounting medium containing DAPI (#H-1200-10, Vector Laboratories) and covered with glass coverslips. Sections were imaged with a wide-field fluorescence microscope (BZ-X810, Keyence). For electrophysiology, the fluorescence came from the DiI that was coated on the Neuropixels probe. The DiI-labeled tracks in coronal sections were aligned to the Allen Common Coordinate Framework^31^ using the SHARP-TRACK software^77^. Reconstructed probe tracks were visualized within the Allen CCF using Brainrender^78^, providing anatomical positioning of the recording sites.

### Chemogenetics: Histology, imaging, and cell detection

For the chemogenetic experiments, the brain was sectioned at a thickness of 40 µm, and all consecutive slices were collected to ensure that capturing the target regions for analysis. During imaging, brain slices containing the target regions (RSP, AIv, TT, and VPM) were selected based on atlas alignment to maintain consistency across samples. For each brain region, 3 consecutive slices were imaged, using a 20x objective lens with a wide-field fluorescence microscope (BZ-X810, Keyence). The images were analyzed using the QUINT workflow^79^, with brain atlas registration based on the Allen Mouse Brain Atlas Common Coordinate Framework version 3 (CCFv3, 2017). The QUINT workflow was structured as follows: first, the images were pre-processed using Nutil^80^ to adjust file names according to the QUINT convention and to downscale as needed for compatibility. Following this, registration to the atlas was performed in two stages. QuickNII^81^ facilitated initial linear alignment, and VisuAlign was subsequently employed to refine the registration through non-linear adjustments. Once registered, specific image features were extracted using ilastik^82^. Quantification of these features was then conducted in Nutil. Since whole-brain imaging was not performed, the “Proportion of input (%)” refers to the proportion of input cells within the imaged brain sections rather than the entire brain.

### Estimates for the number of PT, IT, PT^Fezf2^, and IT^PlxinD1^ neurons

To estimate the number of PT and IT neurons in the medial frontal cortex, we extracted cell counts from the “MERFISH-C57BL6J-638850 with Imputed Genes + Reconstructed Coordinates” data set^83^, which was publicly available in the Allen Brain Cell Types Knowledge Portal (https://knowledge.brain-map.org/abcatlas; “Allen Mouse Common Coordinate Framework, 2020 release”). We restricted the analyses to the ACAd and MOs regions of the isocortex, including their laminar subdivisions (e.g., ACAd1, ACAd2/3, ACAd5, ACAd6a, and ACAd6b). For each region, we classified cells as neuronal (Glut, GABA, and Glut–GABA) or non-neuronal (NA). We further classified the glutamatergic neurons into subpopulations of interest: ET (022 L5 ET CTX Glut), CT (030 L6 CT CTX Glut), or IT (004 L6 IT CTX Glut, 005 L5 IT CTX Glut, 006 L4/5 IT CTX Glut, and 007 L2/3 IT CTX Glut). Cell counts for the different classifications were tabulated for presentation.

Although a prior study showed that the *Fezf2-2A-CreER* and *PlexinD1-2A-CreER* animals capture selectively cells belonging to the PT and IT subtypes of excitatory neurons^25^, it is unclear if the mice can be used to effectively express transgenes in all cells belonging to those subpopulations. To estimate the number of PT^Fezf2^ and IT^PlexinD1^ neurons in the medial frontal cortex, we generated *Fezf2-2A-CreER;Ai75* or *PlexinD1-2A-CreER;Ai75* mice by crossing the respective inducible Cre-driver mice with the *Ai75* reporter mouse for Cre-dependent expression of tdTomato in the nucleus. At 3–4 weeks of age, each animal received a unilateral stereotaxic injection of AAV9-mDlx-GFP-Fishell-1 into the ACAd/medial MOs for GFP expression in GABAergic neurons. One week later, tamoxifen was administered daily for five consecutive days to induce tdTomato labeling of PT^Fezf2^ or IT^PlxnD1^ neurons. Three weeks after the final tamoxifen injection, mice were deeply anaesthetized, perfused with ice-cold PBS followed by 4% paraformaldehyde in PBS, and brains were post-fixed overnight at 4 °C. Coronal sections were cut at 40 μm thickness using a vibratome. Free-floating sections were washed twice in PBS, permeabilized for 10 min in 0.3 % Triton X-100/PBS (room temperature, protected from light), rinsed twice in PBS, and blocked for 1 h in blocking solution (5 % normal serum; Abcam AB7475, 0.1 % Triton X-100 in PBS). Sections were incubated overnight at 4 °C with mouse anti-NeuN (#MAB377, Millipore) primary antibody (diluted in blocking solution), washed three times in PBS (10 min each), and incubated for 1 h at room temperature with Alexa Fluor 405-conjugated secondary antibody (diluted in the same blocking solution). After three final PBS washes, sections were mounted with VECTASHIELD HardSet Antifade Mounting Medium without DAPI (#H-1400-10, Vector Laboratories). Images were acquired on a Zeiss LSM 880 confocal microscope. The images were analyzed in Fiji/ImageJ to obtain cell counts. Specifically, we quantified the proportion of excitatory neurons (i.e., those that are NeuN-positive but GFP-negative) that expressed tdTomato, indicating that they were either PT^Fezf2^ or IT^PlxnD1^ neurons.

### Two-photon imaging: Imaging procedures

Two-photon imaging was performed using a Movable Objective Microscope (MOM, Sutter Instrument) equipped with a resonant-galvo scanner (Rapid Multi Region Scanner, Vidrio Technologies) and a water-immersion 20X objective (XLUMPLFLN, 20x/1.00 N.A., Olympus). Image acquisition was controlled by ScanImage 2020 (Vidrio Technologies). To visualize dendrites of PT^Fezf2^ neurons labeled with EGFP and axonal boutons labeled by Synaptophysin-mRuby2, two femtosecond-pulsed lasers were aligned to converge into the excitation path: a fixed-wavelength laser for excitation at 920 nm (Axon 920-2, Coherent) and a fixed-wavelength laser for excitation at 1064 nm (ALCOR 1064-2 W, SPARK Lasers). Emitted fluorescence signals were separated using dichroic mirrors and collected through a 475–550 nm bandpass filter for EGFP and a 580–680 nm bandpass filter for mRuby2. Power after the objective was typically below 20 mW for each laser. For longitudinal imaging of the same field of view, power levels for the lasers were held constant across sessions to ensure consistency.

During each imaging session, the mouse was head-fixed under 0.8-1% isoflurane anesthesia delivered via a nose cone. Body temperature was maintained at 37.4°C using a feedback-controlled heating pad system (FHC #40-90-8D). Imaging was targeted to the ACAd/medial MOs region within 400 μm of the midline. Apical tuft dendrites were identified within 0–200 μm of the pial surface, and segments located at 20–100 μm depth were selected for imaging. For each field of view, a z-stack was acquired for a depth of between 10–30 μm and at 0.5-μm intervals using bidirectional scanning at 15 Hz, with 1024 × 1024 pixel resolution and a spatial resolution of 0.094 μm per pixel. The same mouse would be imaged repeatedly at the same fields of view on days −3, −1, 1, and 3 relative to drug administration. No imaging was performed on day 0. On that day, the animal received an intraperitoneal injection of either psilocybin (1 mg/kg) or saline (10 mL/kg) while awake. Following the injection, the mouse would be transferred to a clean cage for observation of head twitch responses for 10 minutes before being returned to their home cage.

### Two-photon imaging: Data analysis

Motion correction of the two-photon imaging data was performed using the MultiStackReg plugin in Fiji, with the transformation matrix derived from the green channel applied identically to the red channel. To quantify the change across sessions, the z-stack images for a field of view for the four time points (days −3, −1, 1, and 3) were imported. Based on the brighter EGFP signals in the green channel, we selected a narrower z-range that was consistently captured across days and converted to single images via maximal projection. Next, using custom MATLAB scripts, for each channel of each image, we subtracted the background, which was estimated by selecting a small region of interest in each image devoid of any neural structure to determine the mean background intensity value. Any pixel with a negative value after subtraction was set to zero. To equalize brightness across sessions, we then clipped pixel values to a range (10–160), and then linearly stretched them back to the full range (0–255). To further correct for potential misalignments, images were rigidly registered, using the green channel of the image from the first time point as reference via MATLAB’s “imregtform” function (rigid, multimodal metric). We cropped the images to isolate a portion of the field of view that can be consistently observed across all time points.

Initially, by visualizing the EGFP-expressing dendrites of PT^Fezf2^ neurons and mRuby2-expressing axonal boutons in the same field of view, our goal was to identify specifically those axonal boutons from RSP that apposed dendritic spines in the medial frontal cortex. The mRuby2 was fused to synaptophysin, which worked well to localize the fluorophore selectively to axonal boutons. However, the approach resulted in dense labeling of PT^Fezf2^ neurons, which made the dual-color analysis unreliable due to the many overlapping dendrites and therefore difficulty in characterizing all the spines along the dendrites. For this reason, we focused the analysis solely on the mRuby2-expressing axonal boutons.

We developed custom MATLAB scripts for automated detection of red fluorescent puncta. From the red channel of the image, we generated a binary mask by detecting pixel values that lied above an intensity threshold and requiring the pixel value to exceed the value at the same location in the green channel. The same threshold was used for analyzing all images and was chosen based on inspecting the histograms of pixel values from many images. Puncta (i.e., connected pixels in the binary mask) were detected, and puncta that were too small (<5 pixels) or too large (>1000 pixels) were excluded from further analysis. The number of puncta was determined for each image. Change in the number of puncta across sessions were normalized as percent change relative to the baseline, which was defined as the average number of puncta observed in day -3 and -1.

### Slice electrophysiology: Sample preparation

The aim was to label PT^Fezf2^ neurons with a red fluorophore for targeted whole-cell recording, while expressing ChR2 in the axons of RSP→ACAd neurons for photostimulation. We used two different strategies to label PT^Fezf2^ neurons for slice electrophysiology. For the first strategy, we generated *Fezf2-2A-CreER;Ai14* mice to label PT^Fezf2^ neurons, and injected AAV9-hSyn-hChR2(H134R)-EYFP (#26973, Addgene) into RSP to express ChR2. This was our preferred strategy because PT^Fezf2^ neurons had bright and uniform tdTomato expression. Due to limited availability of the double transgenic mice in our colony, we also used a second strategy: using *Fezf2-2A-CreER* mice, AAV1-FLEX-tdTomato (#28306, Addgene) was injected into ACAd/medial MOs to label PT^Fezf2^ neurons, and AAV9-hSyn-hChR2(H134R)-EYFP (#26973, Addgene) was injected into RSP to express ChR2. For both strategies, 3- to 4-week-old animals underwent stereotaxic surgeries for viral injection. Tamoxifen was administered daily for five consecutive days to induce tdTomato labeling of PT^Fezf2^ neurons. Then at 7-8 weeks of age, brain slices were prepared from these mice for electrophysiological recordings.

Mice were randomly assigned to receive either psilocybin (1 mg/kg, i.p.) or saline (10 mL/kg, i.p.). Slice electrophysiology experiments commenced either 24 hours or 3 days after drug administration. On the day of the experiment, mice were anesthetized with isoflurane and perfused intracardially with an ice-cold cutting solution containing (in mM): 110 choline chloride (#C1879, Sigma-Aldrich), 25 NaHCO_3_(#S6014, Sigma-Aldrich), 2.5 KCl, 7 MgCl_2_, 1.25 NaH_2_PO_4_(#71505, Sigma-Aldrich), 0.5 CaCl_2_, 11.6 sodium ascorbate (#11140, Sigma-Aldrich), 3.1 sodium pyruvate (#P2256, Sigma-Aldrich), and 20 glucose (#G8270, Sigma-Aldrich). Acute coronal brain slices (300 µm) containing the ACAd/ medial MOs were prepared using a vibratome (VT1000S, Leica Biosystems). The chamber in the vibratome was filled with the cutting solution and surrounded by ice. Following cutting, slices were transferred to artificial cerebrospinal fluid (aCSF) containing (in mM): 127 NaCl, 25 NaHCO_3_, 2.5 KCl, 2 CaCl_2_, 1.25 NaH_2_PO_4_, 1 MgCl_2_, 1 sodium pyruvate, and 20 glucose. Slices were incubated at 34°C for 30 minutes, then maintained at room temperature for at least another 30 minutes before recording. Cutting solution and aCSF were prepared using deionized filtered water (18.2 MΩ·cm) and passed through 0.22 µm filters. The cutting solution and aCSF were continuously bubbled with 95% O_2_ and 5% CO_2_ for at least 15 minutes before use and throughout the slicing and recording procedures.

### Slice electrophysiology: Recording procedure

Slices were transferred to a submerged recording chamber and continuously perfused with oxygenated aCSF (2–3 mL/min) at 30–32°C. Whole-cell voltage-clamp recordings were performed, as previously described^15^, on pyramidal neurons located in layer 5 of ACAd and medial MOs, identified by infrared-differential interference contrast and fluorescence. PT^Fezf2^ neurons were identified by their red fluorescence, while non-PT^Fezf2^ neurons were identified by the absence of fluorescence and their morphology (i.e., a triangular somata and a prominent apical dendrite). We recorded sequentially from adjacent PT^Fezf2^ and non-PT^Fezf2^ neurons that lie at the same depth, typically 30–50 μm below the slice surface, and within 50 μm apart laterally. Although the PT^Fezf2^ neuron was typically recorded first, we have also recorded the non-PT^Fezf2^ neuron first in some cases. Recording pipettes were pulled from borosilicate glass capillaries (BF-150-86-10, Sutter Instruments) using a horizontal puller (P-97, Sutter Instruments) to a tip resistance of 2–4 MΩ. Pipettes were filled with a cesium-based internal solution containing (in mM): 125 Cs-methanesulfonate (#C1426, Sigma-Aldrich), 8 NaCl (#71376, Sigma-Aldrich), 10 HEPES, 0.5 EGTA (#E4378, Sigma-Aldrich), 2 MgCl_2_, 4 Mg-ATP (#A9187, Sigma-Aldrich), 0.4 Na-GTP (#G8877, Sigma-Aldrich), 10 Na-phosphocreatine (#P7936, Sigma-Aldrich), and 1 sodium ascorbate (pH 7.25–7.3 adjusted with CsOH (#213601000, Thermo Fisher Scientific)). All internal solutions were double filtered through a 0.22 μm syringe filter prior to use. Signals were acquired with a MultiClamp 700B amplifier (Molecular Devices), digitized at 10 kHz using a Digidata 1550A interface and pClamp software (Molecular Devices), and filtered at 2 kHz for voltage-clamp recording. Initial series resistance was <20 MΩ, and cells were excluded if access resistance exceeded 25 MΩ at the end of the recording.

Photostimulation of the ChR2-expressing RSP axons was achieved via blue LED illumination (470 nm, Thorlabs). Light pulse (2 ms duration) was delivered through a 40× water-immersion objective (0.8 NA, Olympus). To isolate optically evoked excitatory postsynaptic currents (oEPSCs), cells were held at –70 mV. To isolate optically evoked inhibitory postsynaptic currents (oIPSCs), cells were held at 0 mV. In some experiments, 1 μM tetrodotoxin (TTX, #1078/1, Tocris Bioscience) and 100 μM 4-aminopyridine (4-AP, #0940/100, Tocris Bioscience) were applied to block action potentials and restore synaptic release by enhancing terminal depolarization, respectively. LED intensity was typically kept within ∼0.2–2.0 mW/mm² to avoid triggering network bursts or abnormal oEPSCs. A higher-intensity pulse was occasionally applied to determine the maximal synaptic response in each neuron when needed. For all paired recordings, the light stimulation intensity was kept constant to ensure comparability of synaptic input strength across the two cells. In the recordings performed in the presence of TTX and 4-AP to isolate monosynaptic responses, higher light intensities (compared to recordings without TTX and 4-AP) were used. This adjustment was necessary because synaptic transmission under TTX requires stronger depolarization of presynaptic terminals, and network activity is suppressed by TTX, allowing for the use of stronger stimulation without inducing polysynaptic activity. To examine presynaptic function, paired-pulse ratio (PPR) was assessed using two consecutive light pulses (2 ms duration) with a 100 ms inter-pulse interval.

### Slice electrophysiology: Data analysis

Electrophysiological data were analyzed using Clampfit 10.4 (Molecular Devices). For each type of recording, 5–15 consecutive trials with 15 s inter-trial interval were collected. Traces were aligned to stimulus onset and trial-averaged to obtain the response per cell for each measurement. Peak amplitudes of oEPSCs and oIPSCs were determined from the trial-averaged trace for each cell relative to its baseline (10 ms prior to stimulus). PPR was calculated as the ratio of the second oEPSC peak amplitude to the first oEPSC peak amplitude (oEPSC_2_ / oEPSC_1_) evoked by the two consecutive light pulses. Individual data points represent single recorded neurons or sequentially recorded cell pairs.

## QUANTIFICATION AND STATISTICAL ANALYSIS

### Statistical analysis

Analyses were performed in GraphPad Prism 10.0, MATLAB, Python, and R.

For the network selectivity analysis of the rabies tracing data, we used the chi-squared test as described in an earlier section.

We fit linear mixed effects models to analyze datasets with hierarchical structure (e.g. multiple cells recorded from the same mouse, multiple brain sections originating from the same animal), implemented using the “lmer” function in the lme4 package in R. For two-photon imaging data, we used a linear mixed effects model with fixed effects terms of drug (saline or psilocybin), time (day -3, -1, 1, or 3) and their interaction, with fields of view per mouse modeled as nested random intercepts to account for repeated measures. For slice electrophysiology, dependent measures oEPSC amplitudes and PPR were each fit with linear mixed effects models using fixed effects terms of drug (saline or psilocybin), time (24 hr or 3 day), cell type (PT^Fezf2^ or non-PT^Fezf2^) and all interactions, with cell pairs per brain slice per mouse modeled as nested random intercepts. For slice electrophysiology PT^Fezf2^/non-PT^Fezf2^ ratios, we used linear mixed effects models with fixed effects terms of drug (saline or psilocybin), time (24 hr or 3 day) and their interaction, with random intercepts for brain slices per mouse. For Neuropixels single-unit data, we used a linear mixed effects model with fixed effects terms of drug (saline or psilocybin), time (pre- or post-drug), cell type (opto-tagged or untagged), and all interactions, with cells per mouse modeled as nested random intercepts. For Neuropixels LFP data, we used a linear mixed effects model with fixed effects terms of drug (saline or psilocybin), frequency (slow, delta, theta, beta, or gamma) and their interaction, with a random intercept for each mouse. For chemogenetics data, we evaluated region-specific effects by fitting separate linear mixed effects models for each brain region. Each model included fixed effects of drug (saline or psilocybin), chemogenetic manipulation (DMSO or DCZ), and their interaction, with a random intercept for each mouse. When the linear mixed effects model indicated significant main or interaction effects, we performed post hoc pairwise comparison with Bonferroni correction via estimated marginal means using the emmeans package in R.

For comparisons between two sample groups, when per-group sample size was <10, we used the two-sided Wilcoxon rank-sum test, such as for testing the total number of starter or input cells in the whole brain. For comparisons between two sample groups, when per-group sample size was ≥ 10, we used the two-sided t-test, such as for comparing the baseline firing rates for opto-tagged cells between saline and psilocybin condition, or the paired t-test for paired data. For comparisons of cumulative distributions, we used two-sample two-sided Kolmogorov-Smirnov test, such as for comparing the time to first spike after laser onset for opto-tagged cells between saline and psilocybin condition.

Mean values are presented as mean ± SEM. Statistical significance was defined as *P* < 0.05.

