## Supplementary Figures for "Psilocybin triggers an activity-dependent rewiring of large-scale cortical networks"

**Supplemental Figures 1 – 7**

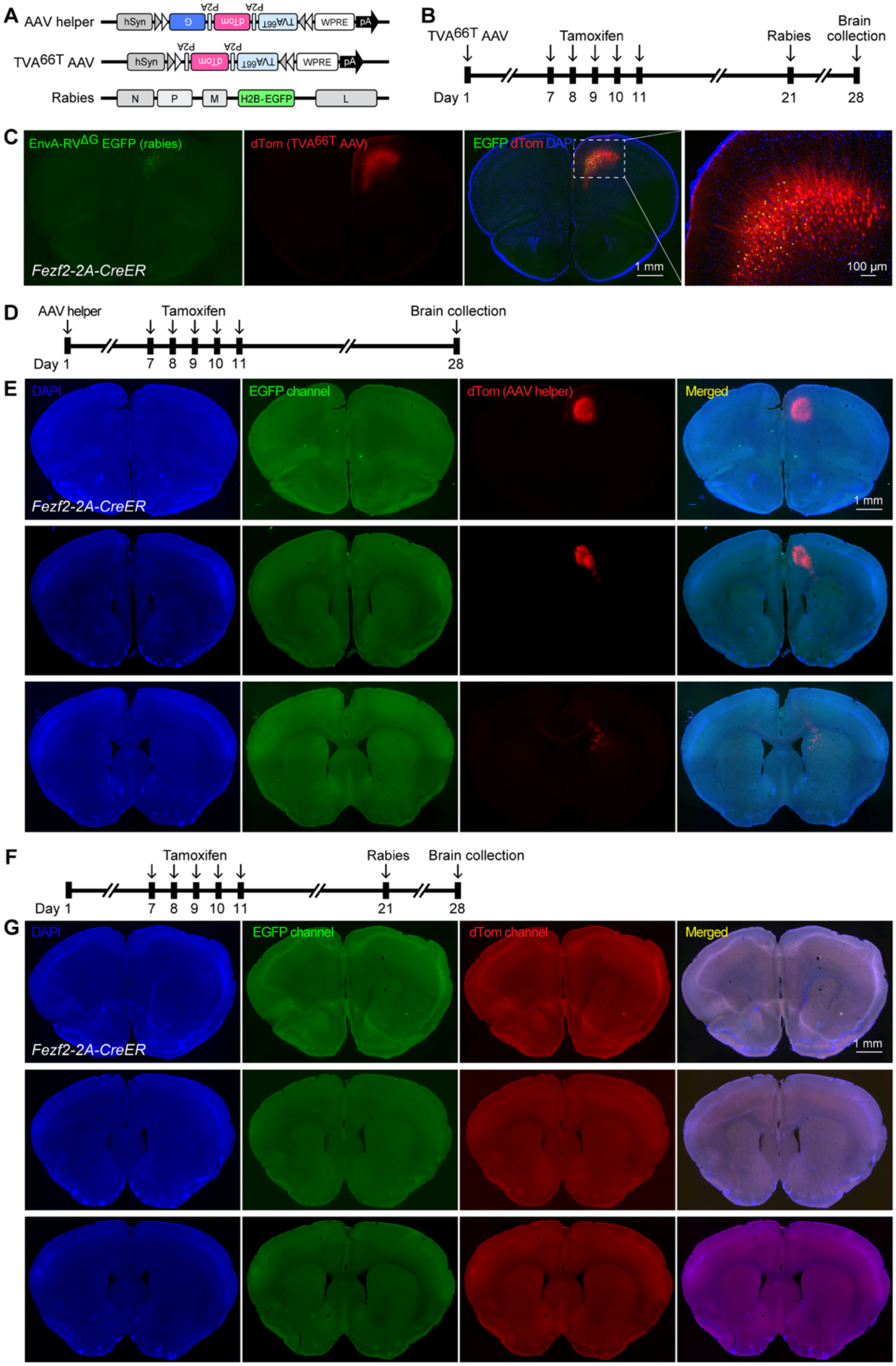


**Figure S1. Control experiments validating monosynaptic tracing in *Fezf2-2A-CreER* mice, related to Figure 1.**

**(A)** Schematic of viral vectors used: Cre-dependent AAV helper (AAV1-hSyn-DIO-TVA$^{66T}$-dTomato-CVS N2c G), Cre-dependent AAV control without G protein (AAV1-hSyn-DIO-TVA$^{66T}$-dTomato), and EnvA-pseudotyped G-deleted rabies (EnvA-CVS N2c$^{\Delta G}$-H2B-EGFP).

**(B) Experiment involving the Cre-dependent AAV control and G-deleted rabies virus.** Experimental timeline.

**(C)** Representative images showing EGFP (green) and dTomato (red) expression restricted to the injection site around the medial frontal cortex of a *Fezf2-2A-CreER* mouse, with nuclear staining by DAPI (blue). As expected, the G-deleted rabies virus could not spread due to the lack of G protein.

**(D) Experiment involving the Cre-dependent AAV virus only.** Experimental timeline.

**(E)** Representative images showing dTomato expression (red) at the injection site in the medial frontal cortex of a *Fezf2-2A-CreER* mouse brain. As expected, there was no green fluorescence because the G-deleted rabies virus was not injected.

**(F) Experiment involving the G-deleted rabies virus only.** Experimental timeline.

**(G)** Representative images showing sections corresponding to the medial frontal cortex of a *Fezf2-2A-CreER* mouse. As expected, the G-deleted rabies virus could not enter any cell because the neurons lack the TVA receptor.

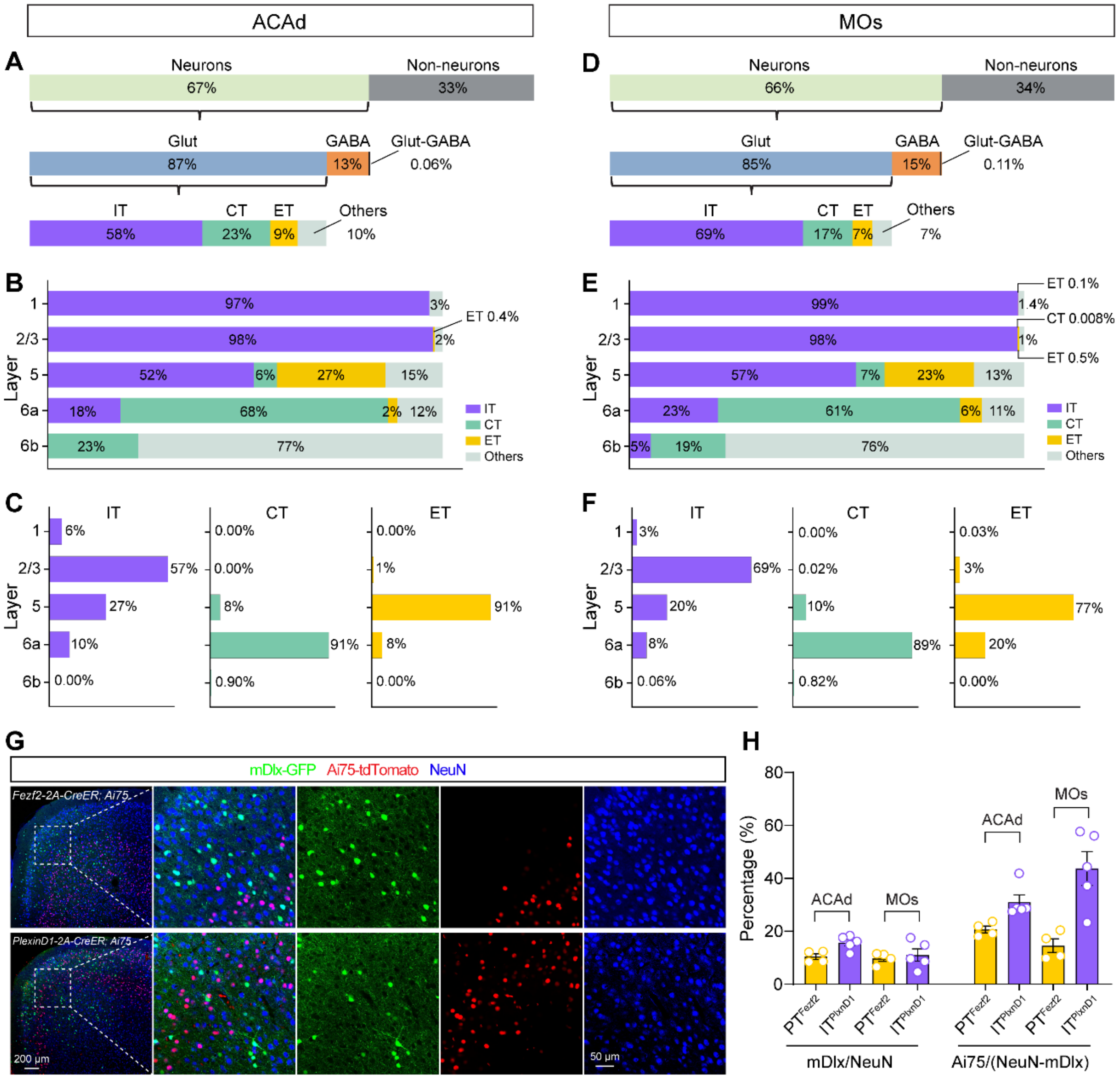


**Figure S2. Estimates for the proportion of excitatory cells as PT, IT, PT$^{Fezf2}$, and IT$^{PlxnD1}$ neurons in the mouse dorsal medial frontal cortex, related to Figure 1, 2.**

**(A)** Proportion of cell types extracted from the publicly available spatial transcriptomics data hosted by Allen Institute (Yao Z et al., Nature, 2023), focusing on the ~14,900 glutamatergic (Glut) neurons in ACAd. IT, intratelencephalic. CT, corticothalamic. ET, extratelencephalic.

**(B)** Proportion of layer 1, 2/3, 5, 6a, and 6b neurons that are IT, CT, ET, or other glutamatergic neurons in ACAd.

**(C)** Proportion of IT, CT, and ET neurons that reside in layer 1, 2/3, 5, 6a, or 6b of ACAd.

**(D-F)** Similar to (A-C) for MOs.

**(G)** AAV-mDlx-GFP was injected into the ACAd/medial MOs region of a *Fezf2-CreER;Ai75* mouse, therefore GFP was expressed in GABAergic cells, tdTomato was expressed in PT$^{Fezf2}$ neurons, and NeuN in blue was used to stain all neurons (top row). Similar strategy was applied to a *PlxnD1-CreER;Ai75* mouse, therefore GFP was expressed in GABAergic cells, tdTomato was expressed in $IT^{PlxnD1}$ neurons, and NeuN in blue was used to stain all neurons (bottom row). Images from fixed coronal sections showing the dorsal medial frontal cortex.

**(H)** Quantification of the fraction of all neurons that were GABAergic in ACAd and medial MOs (left) and estimated fractions of excitatory (non-GABAergic) neurons that were $PT^{Fezf2}$ or $IT^{PlxnD1}$ neurons (right).

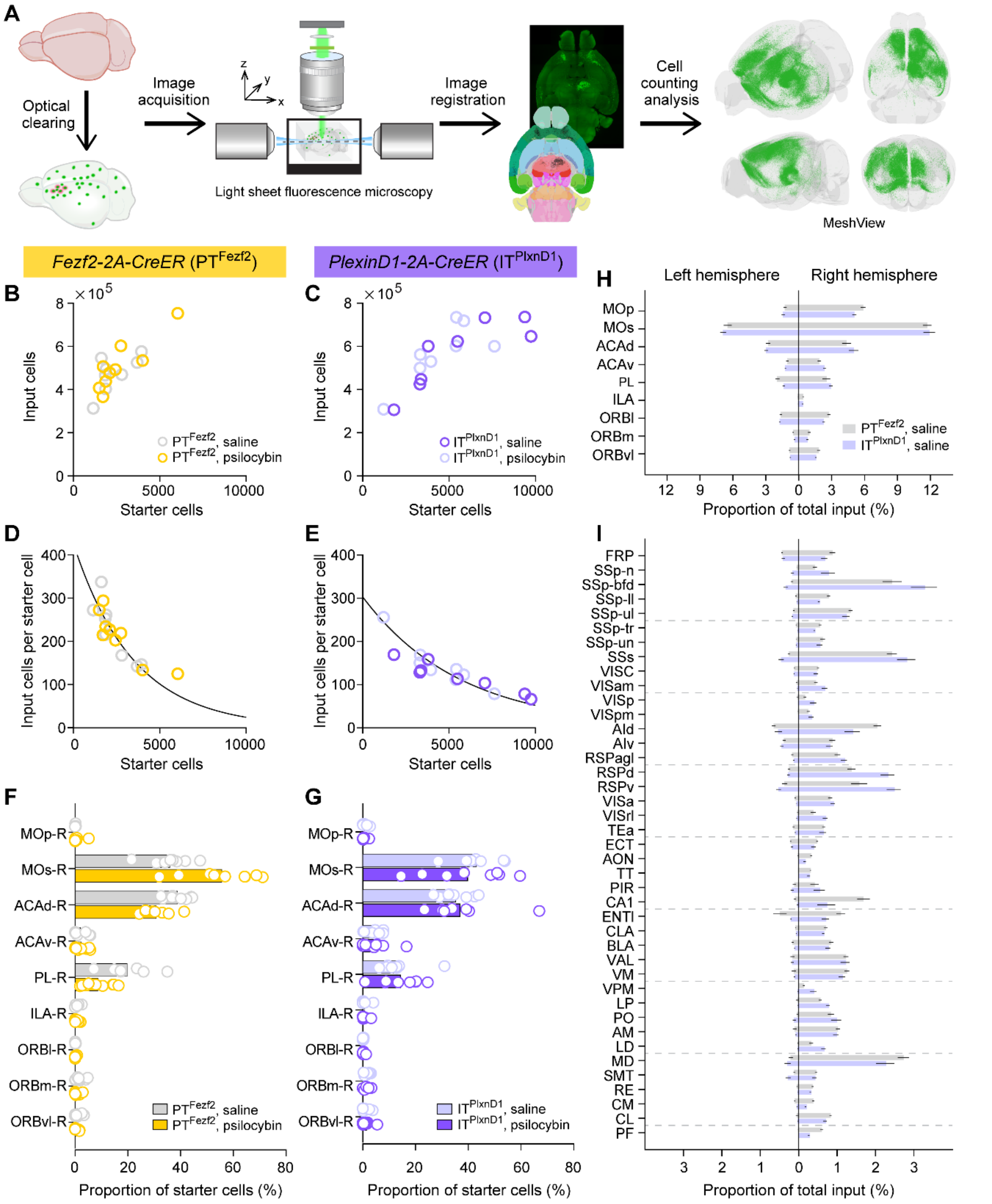


**Figure S3. The distribution of input and starter cells in PT$^{Fezf2}$ and IT$^{Plxnd1}$ mice after saline or psilocybin treatment, related to Figure 1, 2.**

**(A)** Workflow for whole-brain imaging and analysis. Following tissue clearing, brains were imaged using a light sheet fluorescence microscope. The images were registered to the Allen Brain Atlas for cell counting analysis. The EGFP-expressing input cells were visualized in MeshView.

**(B)** Relationship between the total number of input and starter cells in the brain in $PT^{Fezf2}$ mice. Open circle, individual animal.
**(C)** Similar to (A) for $IT^{PlxnD1}$ mice.
**(D)** The number of input cells per starter cell as a function of the number of starter cells in the brain of $PT^{Fezf2}$ mice. Line, exponential decay fit to data from the saline condition. Open circle, individual animal.
**(E)** Similar to (C) for $IT^{PlxnD1}$ mice.
**(F)** The distribution of starter cells in various frontal cortical regions in $PT^{Fezf2}$ mice for the saline and psilocybin groups. Bar, mean. Open circle, individual animal.
**(G)** Similar to (E) for $IT^{PlxnD1}$ mice.
**(H)** Proportion of total input cells in the left and right hemispheres for frontal cortical regions of $PT^{Fezf2}$ and $IT^{PlxnD1}$ mice in the saline condition (mean ± SEM).
**(I)** Similar to (G) for all other regions in the brain.

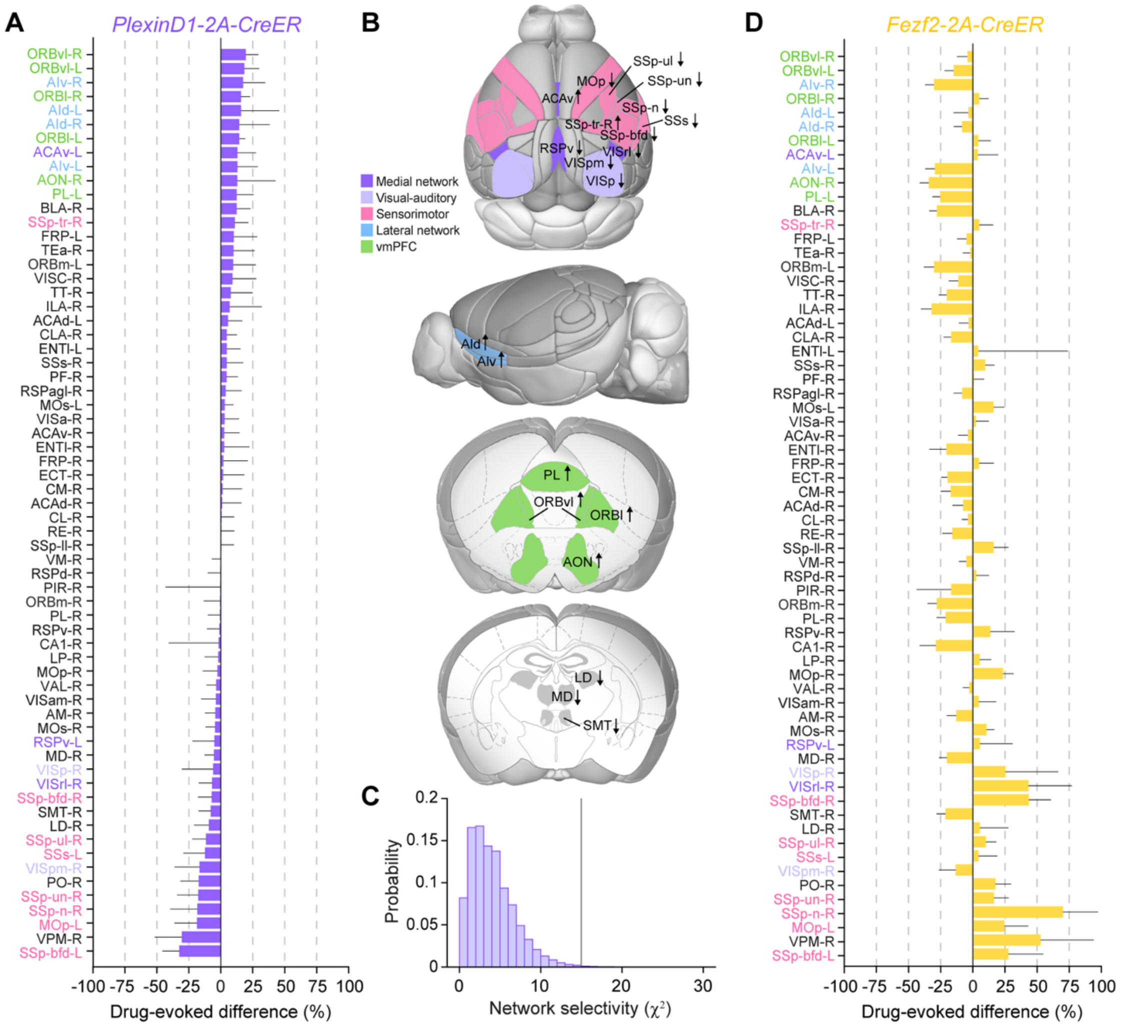


**Figure S4. Drug-evoked difference and network selectivity for IT neurons in *PlexinD1-2A-CreER* mice, related to Figure 2.**

**(A)** Drug-evoked difference (psilocybin subtracted by saline, relative to saline) in the proportion of input cells for all 65 presynaptic regions for $IT^{PlxnD1}$ neurons (mean and 90% confidence interval). The list of presynaptic regions was sorted based on the drug-evoked difference value for $IT^{PlxnD1}$ neurons. Presynaptic regions with >10% or <-5% in drug-evoked difference were color-coded according to their network membership.

**(B)** Schematic showing the location of the presynaptic regions with >10% or <-5% in drug-evoked difference for $IT^{PlxnD1}$ neurons. The presynaptic regions are color-coded according to their network membership.

**(C)** Network selectivity analysis, testing against the null hypothesis that the increases and decreases in drug-evoked difference for $IT^{PlxnD1}$ neurons would be distributed randomly across the 5 cortical networks. Histogram, the distribution of $\chi^2$ values for the null hypothesis. Vertical line, the observed $\chi^2$ value ($P$ = 0.0047).

**(D)** Drug-evoked difference (psilocybin subtracted by saline, relative to saline) in the proportion of input cells for all 65 presynaptic regions for $PT^{Fezf2}$ neurons (mean and 90% confidence interval). The list of presynaptic regions was sorted based on the drug-evoked difference value for $IT^{PlxnD1}$ neurons (i.e., same order as Fig. S4A).

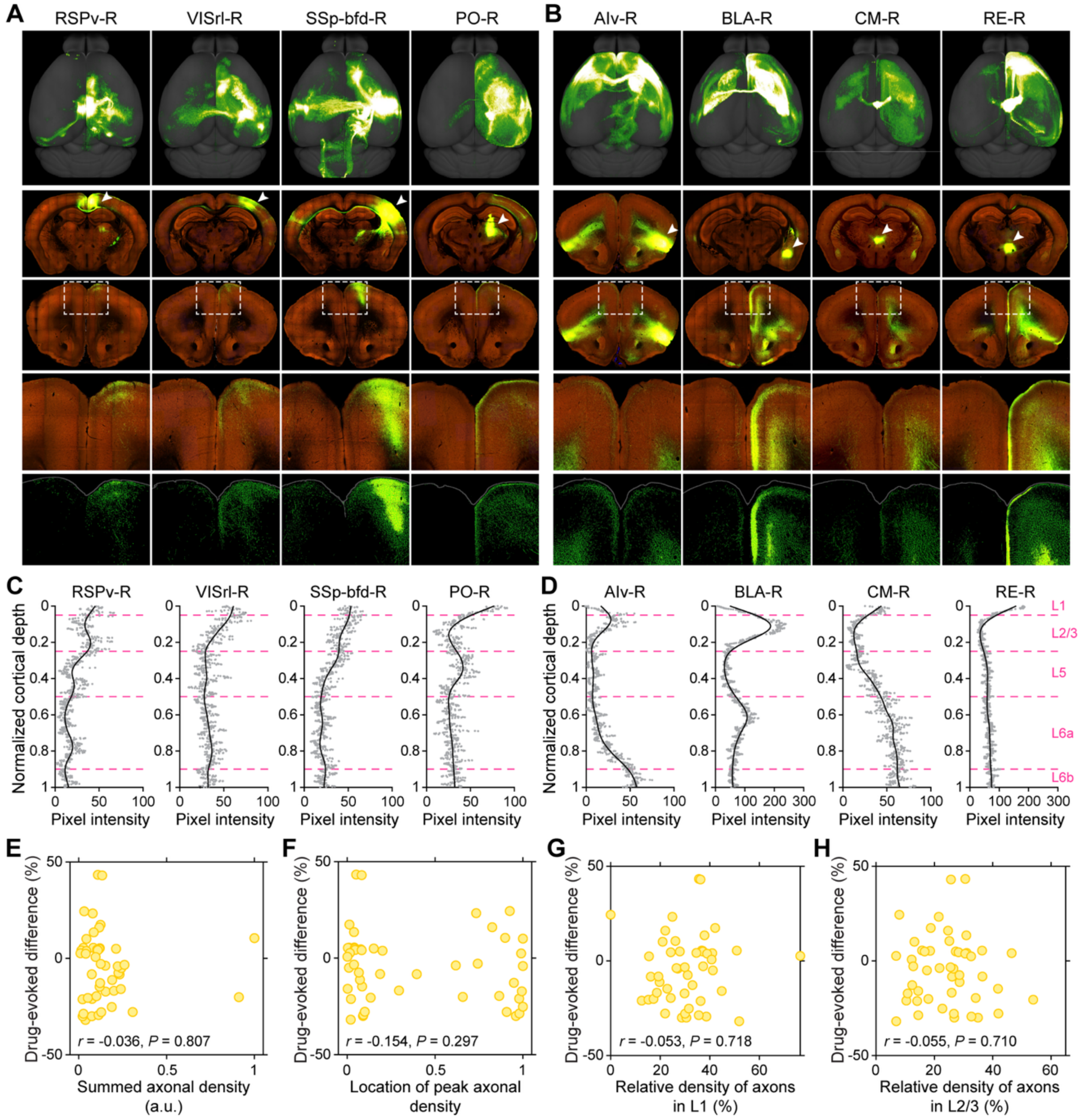


**Figure S5. Analysis of the laminar profile of axonal density from presynaptic regions, related to Figure 3.**

**(A)** Representative images of 4 presynaptic regions with increased numbers of input cells following psilocybin treatment, as identified by monosynaptic rabies tracing. Top row, 3D visualization of brain-wide axonal projection emanating from neurons in the presynaptic region. Second row, serial two-photon image showing the section at the injection site (arrowhead). Third row, serial two-photon image showing the section at the dorsal medial frontal cortex. Fourth and fifth rows, magnified view of the framed area in the third row. Red, autofluorescence. Green, fluorophore.

**(B)** Similar to (A) for 4 presynaptic regions with decreased number of input cells.

**(C)** The laminar profile of pixel intensity (as an estimate of axonal density) in presynaptic regions corresponding to the examples shown in (A). Red dashed lines, positions that divide layer 1, layer 2/3, layer 5, layer 6a, and layer 6b.

**(D)** Similar to (C), corresponding to the presynaptic regions depicted in (B).

**(E)** Scatter plot showing how drug-evoked difference in the number of input cells to $PT^{Fezf2}$ neurons may relate to the summed axonal density, which was calculated by adding up the pixel intensity across the entire profile. Filled circle, individual presynaptic region.

**(F)** Similar to (E) for location of peak axonal density, which was determined by finding the laminar position with the highest pixel intensity.

**(G)** Similar to (E) for relative density of axons in layer 1, which was calculated by dividing the summed axonal density in the portion of the profile demarcated as layer 1 by the summed axonal density across the entire profile.

**(H)** Similar to (G) for layer 2/3.

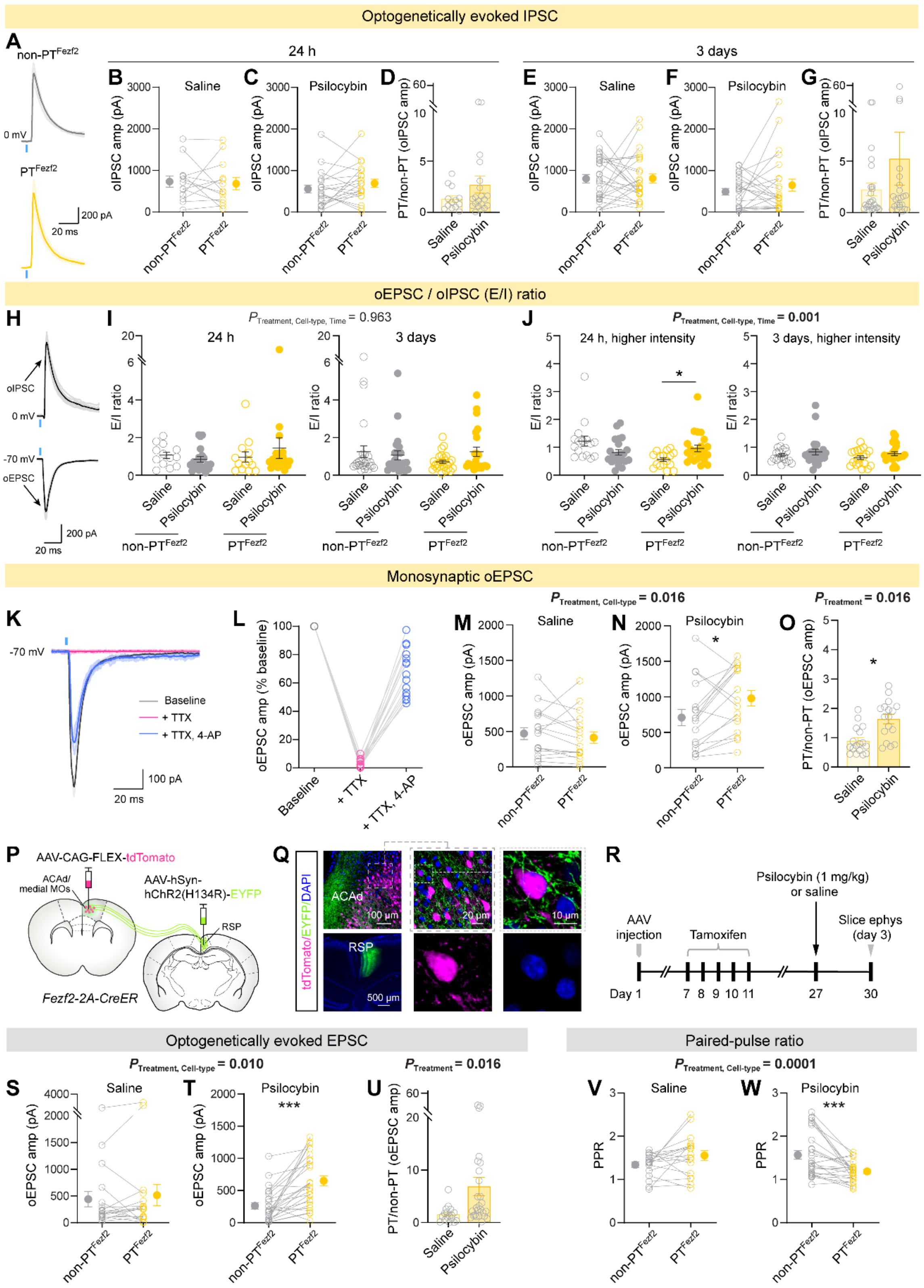


**Figure S6. Additional analyses and data for the slice electrophysiology experiments, related to Figure 4.**

**(A)** Example optogenetically evoked inhibitory postsynaptic current (IPSC) in a pair of $PT^{Fezf2}$ and non-$PT^{Fezf2}$ neurons, for saline and psilocybin condition.
**(B)** Amplitude of the optogenetically evoked IPSC, 24 hr after saline administration. Circle, individual cell.
**(C)** Similar to (B) for psilocybin administration.
**(D)** Based on data in (B) and (C), the ratio calculated by dividing the amplitude of a $PT^{Fezf2}$ neuron by the amplitude of its paired non-$PT^{Fezf2}$ neuron. Circle, individual cell pair. Bar, mean ± SEM.
**(E–G)** Similar to (B–D), 3 days after saline or psilocybin administration.
**(H)** Example oEPSC and oIPSC recorded from the same non-$PT^{Fezf2}$ neuron.
**(I)** Ratio of the oEPSC amplitude divided by the oIPSC amplitude, 24 hr or 3 days after saline or psilocybin administration, for $PT^{Fezf2}$ and non-$PT^{Fezf2}$ neurons. Circle, individual cell.
**(J)** Similar to (I) except using a light pulse with higher intensity. This was because when using standard light intensity, EPSCs were reliably evoked, but IPSCs were sometimes absent. This was why after testing with the standard light intensity, we typically repeated the measurements with a higher light intensity, which could evoke stronger IPSCs. However, sometimes EPSCs have abnormal shape at the higher light intensity, and those cells were excluded from this plot. Considering these caveats, we show both sets of data at the two different light intensities.
**(K)** Example optogenetically evoked EPSC at baseline and after the application of 1 µM TTX or 1 µM TTX and 100 µM 4-AP. TTX and 4-AP were added in order to isolate monosynaptic transmission.
**(L)** Amplitude of the optogenetically evoked EPSC in the different conditions, normalized by baseline. Circle, individual cell.
**(M)** Amplitude of the optogenetically evoked EPSC in the presence of TTX and 4-AP, 24 hr after saline administration. Circle, individual cell.
**(N)** Similar to (M) for psilocybin administration.
**(O)** Based on data in (M) and (N), the ratio calculated by dividing the amplitude of a $PT^{Fezf2}$ neuron by the amplitude of its paired non-$PT^{Fezf2}$ neuron. Circle, individual cell pair. Bar, mean ± SEM.
**(P)** Viral strategy to express ChR2 in RSP neurons. $PT^{Fezf2}$ neurons express tdTomato because AAV-CAG-Flex-tdTomato was injected into the ACAd/medial MOs region of a *Fezf2-2A-CreER* mouse.
**(Q)** Post hoc histology showing the fluorophore expression in medial frontal cortex and RSP.
**(R)** Experimental timeline. Successive whole cell recordings were made from a $PT^{Fezf2}$ neuron and a non-$PT^{Fezf2}$ neuron. A 470 nm LED was used to photostimulate the RSP axons in the acute brain slice.
**(S)** Amplitude of the optogenetically evoked EPSC, 3 days after saline administration. Circle, individual cell.
**(T)** Similar to (S) for psilocybin administration.
**(U)** Based on data in (S) and (T), the ratio calculated by dividing the amplitude of a $PT^{Fezf2}$ neuron by the amplitude of its paired non-$PT^{Fezf2}$ neuron. Circle, individual cell pair. Bar, mean ± SEM.
**(V)** Paired pulse ratio, 3 days after saline administration. Circle, individual cell.
**(W)** Similar to (V) for psilocybin administration.

Data in panels (A – O) came from the same experiments to obtain results shown in Fig. 4E-T (i.e., Ai14-based labeling of $PT^{Fezf2}$ neurons). A – J: N = 13–16 cell pairs from 6 mice for 24 hr after saline, 19–22 cell pairs from 7 mice for 24 hr after psilocybin, 19–26 cell pairs from 5 mice for 3 days after saline, 23–24 cell pairs from 7 mice for 3 days after psilocybin. L: N = 15 cells from 14 animals. M – O: N = 17 cell pairs from 6 mice for 24 hr after saline,

17 cell pairs from 5 mice for 24 hr after psilocybin. Data in panels (P – W) came from the different experiments involving viral-mediated tdTomato expression in frontal cortical $PT^{Fezf2}$ neurons. N = 18 cell pairs from 4 mice for 3 days after saline, 26 cell pairs from 5 mice for 3 days after psilocybin.

For (B, C, E, F), linear mixed effects model with fixed effects terms of drug (saline or psilocybin), time (24 hr or 3 day), cell type ($PT^{Fezf2}$ or non-$PT^{Fezf2}$) and all interactions, with cell pairs per brain slice per mouse modeled as nested random intercepts. For (D, G), linear mixed effects model with fixed effects terms of drug (saline or psilocybin), time (24 hr or 3 day), and interaction, with random intercept for brain slice and animal. Post hoc pairwise comparisons with Bonferroni correction. For (I) and (J), data were analyzed separately using linear mixed-effects models with fixed effects of drug, time, cell type, and all interactions, and random intercepts for cell pair nested within brain slice and animal (Mouse/Slice/CellPair). Post hoc pairwise comparisons between treatment groups were performed for each cell type and timepoint using estimated marginal means with Bonferroni correction. For (M, N), (S, T), (V, W), data were analyzed separately using linear mixed-effects models with fixed effects of drug, cell type, and their interaction, with random intercepts for cell pair nested within brain slice and animal (Mouse/Slice/CellPair). Post hoc pairwise comparisons between cell types were performed within each treatment group using estimated marginal means, with Bonferroni correction. For (O) and (U), the ratio of oEPSC amplitudes (PTFezf2 / non-PTFezf2) was analyzed separately using a model with drug as a fixed effect and brain slice nested within animal as a random intercept (Mouse/Slice). Post hoc comparisons were based on estimated marginal means. See **Table S1**. *, $P < 0.05$. **, $P < 0.01$. ***, $P < 0.001$.

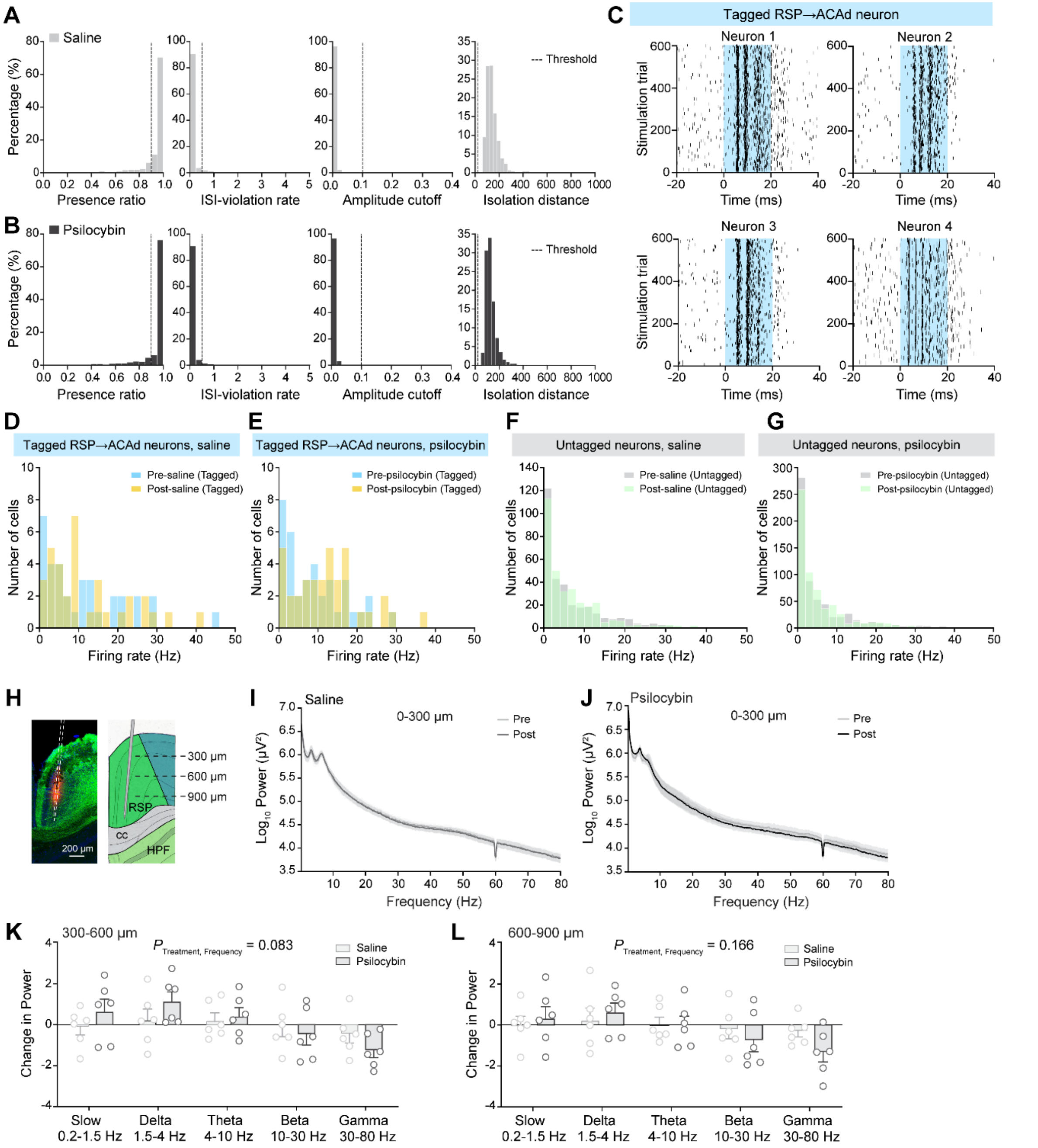


**Figure S7. Additional analyses for the Neuropixels experiments, related to Figure 5.**

**(A)** The distribution of quality metrics for all untagged units in the saline group marked as 'good' after Kilosort and Phy. Dashed line, criterion for the units to be included for further analyses (see Methods).

**(B)** Similar to (A) for the psilocybin group.

**(C)** Additional examples for spike raster plots for tagged RSP→ACAd neurons. Blue, period of laser stimulation.

**(D)** The distribution of firing rates for opto-tagged RSP→ACAd neurons in the pre and post periods around saline administration.

**(E)** Similar to (D) for opto-tagged RSP→ACAd neurons around psilocybin administration.

**(F)** Similar to (D) for untagged RSP neurons.
**(G)** Similar to (D) for untagged RSP neurons around psilocybin administration.
**(H)** This is the same histology figure shown in Figure 5C, delineating rough estimates for depths of 300, 600, and 900 µm.
**(I)** Mean power spectrum calculated from LFP signals from channels within depth of 0 – 300 µm, before and after saline.
**(J)** Similar to (I), before and after psilocybin.
**(K)** Fractional change in spectral power within different frequency after saline or psilocybin, calculated from LFP signals from channels within depth of 300 – 600 µm. Circle, individual animal. Bar, mean ± SEM.
**(L)** Similar to (K), for channels within depth of 600 – 900 µm.

Data came from the same experiments to obtain results shown in Fig. 5. N = 6 mice for psilocybin and 6 mice for saline.

For (K) and (L), linear mixed effects model with fixed effects terms of drug (saline or psilocybin), frequency (slow, delta, alpha, beta, or gamma), and interaction, with a random intercept for animal. See **Table S1**.
